# Population-level migration modeling of North America’s birds through data integration with BirdFlow

**DOI:** 10.1101/2025.09.30.679621

**Authors:** Yangkang Chen, David L. Slager, Ethan Plunkett, Miguel Fuentes, Yuting Deng, Stuart A. Mackenzie, Lucas E. Berrigan, Daniel Fink, Daniel Sheldon, Benjamin M. Van Doren, Adriaan M. Dokter

## Abstract

**Background:** Accurate information on population-level movements of migratory animals is essential for understanding migration and for designing effective conservation strategies in a changing world. Yet such information remains scarce for most migratory species due to the effort and expense needed to collect data across their full distribution ranges. BirdFlow is a probabilistic modeling framework that infers population-level movements from weekly species distribution maps produced by the participatory science project eBird. However, BirdFlow models have only been tuned for a handful of species using high-resolution individual tracking data, which is not available for most migratory species.

**Methods:** Here, we introduce a general tuning and evaluation framework for BirdFlow that enables the first large-scale integration of distributional and individual-level data to infer animal movement across continents and hundreds of migratory species, eliminating reliance on any single individual-tracking data source. By generalizing the BirdFlow model parametrization, we enable tuning and validation using multiple complementary data sources, including GPS tracks, banding recoveries, and radio telemetry data from the Motus Wildlife Tracking System. We investigate the efficacy of this approach by (1) investigating predictive performance compared to null models; (2) validating the biological plausibility of BirdFlow models by comparing movement properties such as route straightness, number of stopovers, and migration speed between model-generated routes and real movement tracks; and (3) comparing the performance of models tuned on species-specific movement data to models tuned using hyperparameters transferred from other species.

**Results:** Our results show that BirdFlow models produced by the new tuning framework achieve biologically realistic performance, even for prediction horizons of thousands of kilometers and several months. When species-specific data are unavailable, models can still be tuned using data from other phylogenetically adjacent species to achieve improved performance.

**Conclusions:** By integrating eBird Status & Trends abundance surfaces with data from banding recaptures, radio telemetry, and GPS tracking, we scale BirdFlow model to 153 North American migratory species, representing the first collection of continental-scale population-level movement and forecasting models. Species-specific tuning improves population-level movement forecasts, while taxonomically informed hyperparameter transfer supports the modeling of data-limited species. Overall, our work offers a foundation for more accurate predictions across hundreds of species for research in ecology and conservation, disease surveillance, aviation, and public outreach.

## Background

Our understanding of continent-scale bird migration has been greatly fostered by modern tracking technologies and the rapid accumulation of participatory science data [1, 2]. Despite these advances, population-level variation in movement behavior remains poorly understood for most bird species, particularly across continental extents and throughout the annual cycle [3].

This gap poses a critical challenge because species’ ranges often span diverse ecoregions and habitats, exposing different populations of the same species to distinct environmental barriers and threats (e.g., [4]). Spatial heterogeneity in threats, which include habitat degradation, artificial light at night (ALAN), climate change, and hunting risk, likely drives corresponding variation in bird population trends [5–10]. Therefore, understanding population-level movements within a species’ range is essential for quantifying migratory connectivity and for targeting conservation actions in a changing world [11]. However, such information remains largely unavailable due to the difficulties of tracking multiple moving populations across species’ range.

Several modeling frameworks have been proposed to infer large-scale avian movement by integrating multiple data types, including methods that combine individual tracking and species distribution information within integrated statistical frameworks [16, 37]. These approaches represent important advances and highlight the growing range of tools available for migration inference at broad spatial scales. However, these approaches remain constrained by the limited availability of individual-level movement data, which limits their application in regions and for species for which tracking data remain sparse.

BirdFlow is a probabilistic modeling framework developed to address this need by inferring the population-level movements across a species’ range from spatiotemporal abundance distributions [12]. The model builds on relative abundance estimates generated by the eBird Status & Trends project (hereafter, eBird S&T) [13, 14] and uses mechanistic optimization that accounts for energy costs and inter-population movement patterns. Performance of the BirdFlow model can be improved when hyperparameters are tuned using individual movement data from other sources, such as GPS and satellite tracking [12]. However, generalizing BirdFlow to hundreds of species has remained challenging because GPS tracking data are unavailable for most migratory birds [15].

To overcome the reliance on any single individual-tracking data source and produce BirdFlow models for many species, we introduce a unified model tuning framework to train BirdFlow models based on general mark-resight data. We also introduce a set of intuitive validation metrics for quantifying the quality and predictive performance of BirdFlow models. Our approach standardizes observations across multiple data sources, including GPS tracking data, banding recovery data, and Motus Wildlife Tracking System automated radio telemetry data (Motus; https://motus.rog; [17]). We address three primary questions: (1) Can integrating multiple data sources enable BirdFlow modeling across hundreds of species? (2) How consistent are optimal model parameterizations among species with similar ecological or morphological traits? and (3) Can such cross-species similarities be leveraged to model species lacking individual-level tracking data? Finally, we discussed how species-specific models tuned using this framework open new opportunities for research and outreach across ecology, evolution, and conservation biology.

## Methods

We describe the tuning and validation framework in the sections below. A schematic summary is presented in Fig. 1.

**Figure 1.**
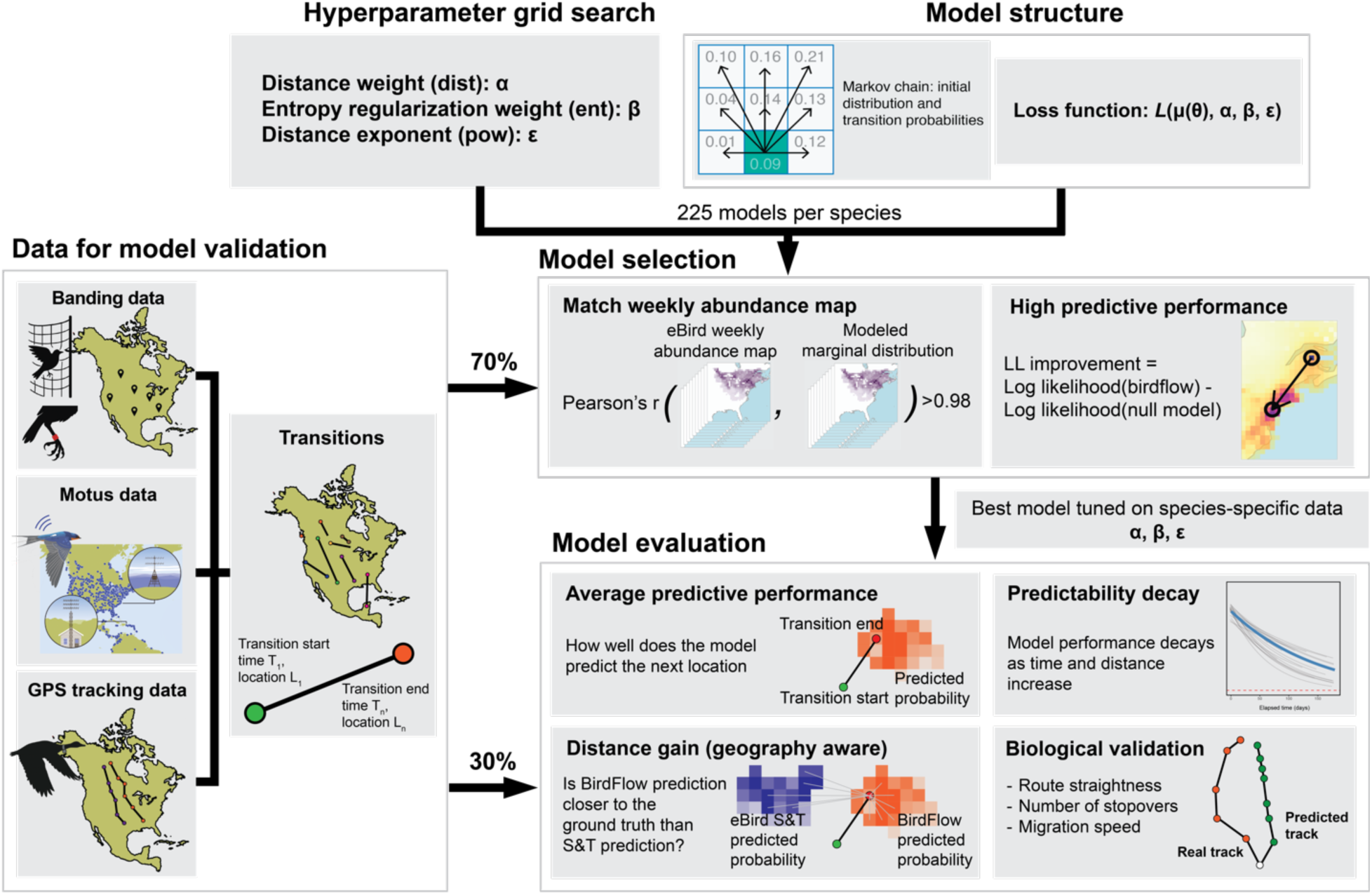
Schematic workflow for the BirdFlow species-specific model tuning and validation framework. The diagram illustrates the main steps of data preparation, model tuning, and validation methods used in this study.

### The BirdFlow model

BirdFlow model structure and training methodology are described in detail in [12]. Briefly, the current version of BirdFlow is a Markovian probabilistic graphical model where each state (location and time) is projected to the next state with a certain transition probability. BirdFlow is trained by optimizing an objective function, with three terms controlled by three hyperparameters:

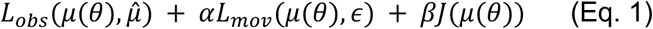

The first term is the “observation” term, which is the same as the “location loss” term in [12]. The observation term minimizes the differences between the modeled probabilistic surface 𝜇(𝜃), given Markov chain parameter set 𝜃 and a species’ spatiotemporal relative abundance surface 𝜇^.

The second term in Eq. 1 is the “distance” term, computed as a function of distance that penalizes longer movements. This requires simulated routes to minimize the movement cost for the population. This term is regulated by hyperparameter 𝛼 (the distance weight) and hyperparameter 𝜖 (the distance exponent). The distance term for a single designated transition is formulated as

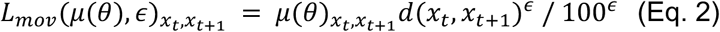

Where 𝜇(𝜃)_*xt,xt*+1_ is the pairwise marginal that specify the probability of moving from cell *x_t_* at time *t* to cell *x*_*t*+1_ at time *t* + 1. *d*(*x_t_*,*x_t+1_*) denotes the great circle distance between *x_t_* and *x*_*t*+1_, with 𝜖 regulating the trade-off of cost between a few long-distance jumps versus many short distance hops. The exponentiated distance d is divided by 100^+^ to make the magnitude of L_mov_ less dependent on 𝜖, thereby reducing correlations in the optimized values of 𝛼 and 𝜖.

The third term in Eq. 1 is the “Shannon entropy” term and is regulated by hyperparameter 𝛽 (the entropy weight), which controls the stochasticity of the routes (Fig. 1).

### Species relative abundance surfaces

We acquired eBird Status & Trends weekly relative abundance distributions (data version 2023; hereafter, S&T abundance surfaces) using the ebirdst package version 3.2023.1 [18]. These surfaces are model-based products derived from eBird checklist data. In the present study, we used these S&T abundance surfaces as inputs to the BirdFlow framework but did not explicitly propagate their uncertainty through the tuning procedure, largely because doing so across many species would be computationally prohibitive. We therefore interpret them as model inputs rather than components of a fully joint uncertainty-propagating inferential pipeline. All species are assigned quality scores by experts during the S&T modeling process; we only included species assigned the highest score (3) across all seasons. We preprocessed S&T surfaces to restrict geographic scope to the Western Hemisphere. In addition, to avoid the influence of high-abundance outliers, we truncated abundance distributions by replacing all values above the 0.99 quantile with the 0.99 quantile value. The rationale behind this trimming was to mitigate the influence of imperfect S&T abundance surfaces on model predictions, where excessive counts in aggregation areas (e.g., due to roosting behavior) could bias abundance estimates. We justified the choice of quantile using a sensitivity analysis (Additional file 1: Fig. S1).

### Individual mark-resight data (transitions)

We used empirical movement data to select the best hyperparameters from a set of 225 candidate models (see Hyperparameter tuning section) for each species. We acquired tracking data from Movebank (movebank.org) and published literature (Additional file 1: Table S1). We acquired bird banding data from the United States Geological Survey (USGS) Bird Banding Program (BBL) database [19]. Motus data were acquired following the Motus Collaboration Policy (https://motus.org/resources/policy) and preprocessed to reduce errors (Additional file 1: Supplementary methods).

We converted all data into species-specific BirdFlow spatiotemporal coordinates using the BirdFlowR package, an R-language interface that implements and operationalizes the BirdFlow model (version 0.1.0.9075) [20]. We refer to a set of detections for a single individual bird movement within a migratory season as a “track”. After compiling empirical movement data, we sampled mark-resight pairs (segments of movement) from the tracks, which we call “transitions”. We sampled up to 10000 transitions for each species, considering the tradeoff of computational cost and model performance, using an iterative sampling procedure (Additional file 1: Supplementary methods). For species to be included, we required availability of at least 20 transitions (justified using a sensitivity analysis; Additional file 1: Fig. S2) and high-quality abundance maps based on a sensitivity analysis, resulting in 153 migratory species in North America spanning 14 orders and 39 families (Additional file 1: Table S2). Across species, the number of training transitions varied from 29 to 7000 (mean = 1673). Banding data accounted for 69.2% of transitions per species on average, and Motus data made up 27.6%. GPS tracking data were available for only seven species, served as the primary data source (> 50% of transitions) for four, and comprised 11.8% of all transitions (Additional file 1: Fig. S3).

### Hyperparameter tuning

For each species, we trained BirdFlow models with different combinations of hyperparameters (𝛼, 𝛽, and 𝜖 introduced in the previous sections) in an exhaustive parameter grid (Additional file 1: Supplementary methods). In total, the training generated 225 models per species. We trained models with a spatial resolution of 150 km and temporal resolution of 1 week. We chose this spatial resolution to balance computation cost while maximizing resolution. The 1-week temporal resolution is inherited from the temporal resolution of the S&T abundance surfaces.

For each species, we used 70% of transitions as the training set for model selection in a hyperparameter grid search. We used two criteria when selecting the best model for a species:

(1) First, we selected models with a high correlation between the modeled probability surface and the eBird S&T relative abundance surface, indicating that the BirdFlow model reproduces an accurate abundance surface. We defined the mean correlation score as the average of the correlation score for pre-breeding migration and post-breeding migration seasons:

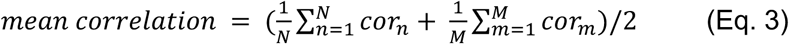

Where 𝑐𝑜𝑟_-_ and 𝑐𝑜𝑟_$_ represent the Pearson correlation coefficient calculated across grid cells between BirdFlow marginal distributions and S&T abundance surfaces at week 𝑛 of pre-breeding migration and week 𝑚 of post-breeding migration, with 𝑁 and 𝑀 being the total number of weeks of pre-breeding migration and post-breeding migration. The mean correlation score thus reflects how well the model projection matches observations during migration, with value 1 representing a perfect match. We only considered models with mean correlation score >0.98, which is an empirical value we chose after investigating the models for tens of species. This threshold ensures any candidate BirdFlow model accurately reproduces a species’ abundance surface across areas of both high and low density.

(2) Second, we selected the best model from the remaining set of candidate models based on how well the model predicted the true resighting location of each transition, quantified by the weighted log-likelihood improvement (𝛥𝐿𝐿). To calculate this metric, we first introduce the log-likelihood improvement (𝛥𝐿𝐿_0_) for a single transition 𝑖. The 𝛥𝐿𝐿_0_ tells us how much more accurate a BirdFlow model prediction is compared to a random sample from the S&T abundance surface for transition 𝑖.

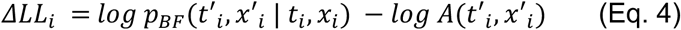

Here, 𝑝_12_ denotes the probability assigned by the BirdFlow model to a transition from a starting location and time to a destination location and time. Specifically, 𝑙𝑜𝑔 𝑝_12_(𝑡′_0_, 𝑥′_0_ | 𝑡_0_, 𝑥_0_) represents the conditional log probability for the BirdFlow model to correctly predict the resighting location 𝑥′ at time 𝑡′, given that the bird is recorded at location 𝑥 at time 𝑡, where [(𝑡_0_, 𝑥_0_), (𝑡′_0_, 𝑥′_0_)] defines empirical transition 𝑖. In simple terms, it describes how probable the observed data are, given the model. 𝑙𝑜𝑔 𝐴(𝑡′_0_, 𝑥′_0_) is the normalized relative abundance at (𝑡′_0_, 𝑥′_0_) calculated from the S&T abundance surface, representing the probability of randomly sampling the exact resighting location from the abundance surface.

The weighted average log-likelihood improvement (𝛥𝐿𝐿) is calculated as

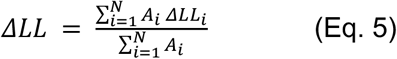

Where 𝑁 represents the total number of transitions and 𝐴_0_ is a weighting factor equal to the normalized weekly relative abundance of the S&T abundance surfaces at the starting location for transition 𝑖. The purpose of the weight is to mitigate potential spatiotemporal bias introduced by rare transitions that are hardly representative of the population.

For each species, we selected the best set of hyperparameters based on the two steps described above. We refer to the models using the selected set of hyperparameters as “species-specifically” tuned models.

### Quantifying hyperparameter transferability

We conducted leave-one-out (LOO) analysis to determine the feasibility of transferring hyperparameters from other species in the scenario that tracking data are not available for a target species. In this process, we sought to determine whether taxonomic information could improve the accuracy of hyperparameter transfer. When applying LOO, we omitted transition data of the target species and selected hyperparameters based on the transition data of the remaining species. We only considered models with a mean correlation score >0.98, consistent with species-specific tuning. We then selected the hyperparameter configuration that had the highest average log-likelihood improvement score within the same taxonomic group as the target species. We compared four model tuning methods, including taxonomically informed (1) family-level LOO (Family-LOO), (2) order-level LOO (Order-LOO), and (3) LOO with all the other species except for the target species (All-LOO). We consider All-LOO as a baseline method since it incorporates no taxonomic information. We also included (4) species-specifically tuned models (no LOO) for comparison.

### Model validation

After tuning, we evaluated tuned models using the remaining 30% of the transition data (test set). We again used log-likelihood improvement to evaluate tuned models as described in the hyperparameter tuning section. However, while log likelihood reflects the quality of prediction, it only validates the model at the exact resighting locations and therefore has limited biologically interpretability. Ideally, an evaluation of a spatiotemporal movement model should also consider how close the predictions are to the resighting locations. To address this need, we introduce a spatially explicit metric called “distance gain” (𝛥𝐷) and a variant, relative distance gain (𝛥𝐷^89:^), as the primary validation metric. It measures how close the BirdFlow model prediction is to the empirical resighting location and compares this distance to a random sample of distances drawn from the abundance surface:

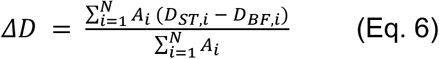

Where 𝐷_12,0_ is the weighted mean distance from the BirdFlow predicted locations to the empirical resighting location for transition 𝑖:

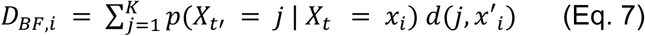

Where 𝑗 is a location index running over all spatial pixels 𝐾. 𝑑(𝑗, 𝑥′_0_) represents the great circle distance between location 𝑗 and the resighting location 𝑥′_0_ of the transition 𝑖. The weight for location 𝑗 is the BirdFlow predicted probability (𝑝) at location 𝑗 time 𝑡′ given the starting location 𝑥_0_ at time 𝑡.

Similarly, 𝐷_BC,0_ is defined as the weighted mean distance from the normalized abundance surface (S&T) to the empirical resighting location for transition 𝑖, representing a “null distance” of drawing a random sample from all potential locations.

To facilitate multi-species comparison, we developed the relative distance gain (𝛥𝐷^89:^ ) to account for the variation of distance gain among species with different migration distances:

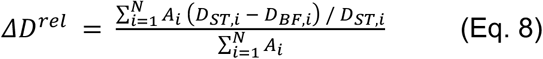

With value 1 representing a perfect prediction of the resighting location, value 0 representing predictive performance equivalent to a random sample from the S&T abundance surfaces, and negative values representing worse performance than a random sample.

### Model decay analysis

Next, we quantified the pattern of decay in model performance across time and distance. We fitted exponential regression functions using the test set transitions and relative distance gain metric. We chose exponential functions because the visual patterns generally match well with exponential decay as transition interval increases, a pattern also seen in [12] (Additional file 1: Supplementary methods). The regression line is asymptotic to y=0, so we define the maximum functional distance horizon as the distance value where the regression line intersects a threshold of 0.05. Thus, the maximum functional distance represents the distance at which BirdFlow model predictions are <5% closer to true resighting locations than a random sample from the S&T distribution of the resighting week (Additional file 1: Supplementary methods, Fig. S4). We do not define a maximum functional time horizon since the models often performed well even beyond the temporal threshold (see results). We also evaluated performance decay using log-likelihood improvement metrics and validated prediction horizons using the same approach.

### Empirical validation of biological metrics

To assess the biological plausibility of BirdFlow-generated trajectories, we compared the route statistics of model-simulated and empirical routes. For all seven species for which GPS tracking data were available at the time of analysis, we extracted track sections during pre-breeding and post-breeding migration phases. Only tracks recorded for more than two weeks and with ≥1 observation per week were included to match the temporal resolution of BirdFlow models. For each track, we sampled 100 trajectory predictions conditional on the starting time and location and the ending time of the track. We then calculated three biological metrics frequently used in migration research [21] for both simulated and observed routes: migration speed, route straightness, and number of stopovers. We defined migration speed as the total distance traveled by the simulated individual within the trajectory divided by the total length of the time.

Route straightness was the beeline distance of the routes divided by the total distance traveled. We defined the number of stopovers as the number of 150-km resolution grid cells in which the individual stayed >7 days; this temporal resolution is predefined and aligned with the eBird S&T product. The route features were calculated using trajr package version 1.5.1 [22] in R version 4.5.

### Modeling hyperparameters using species traits

We modeled the selected hyperparameters for each species using species-specific morphological traits and distribution range statistics to understand hyperparameter variation across species. We acquired species morphological traits from the AVONET dataset [23]. We further calculated range-wide summaries as explanatory variables, including range size, spatial variation of abundance, and the minimum, maximum, and centroid of longitude and latitude for different seasons (Additional file 1: Supplementary methods). Some of these explanatory variables were highly correlated (Pearson’s r >0.8); for a group of correlated variables, we only included one representative variable in the model (Additional file 1: Supplementary methods, Fig. S5). To account for potential overfitting, we tuned Random Forest regression models for each hyperparameter separately using 5-fold cross-validation on the species-level dataset and calculated the coefficient of determination (𝑅^G^) based on pooled predictions on out-of-fold datasets. We calculated impurity-based feature importance metrics for the best model and linearly regressed BirdFlow hyperparameters against the variable with the highest feature importance in the Random Forest model to understand how a single trait is associated with the hyperparameter values. The feature calculations are performed in R version 4.5 and the Random Forest model is fitted using the scikit-learn package version 1.6.1 [24] in Python version 3.11.11.

## Results

### Multi-source transition data improves model performance

BirdFlow models tuned using species-specific mark-resight data consistently predicted resight locations better than a random sample from the S&T abundance surface corresponding to the week of resighting (the null model) for all species (Fig. 2). This was reflected in both relative distance gain and log-likelihood (LL) improvement metrics. Averaged across species and transitions, tuned models were 27 times better in predicting the exact resighting location (average log-likelihood improvement 3.30, range: 0.27–7.13 across species; 95% percentile interval 0.67–5.17) and 1058 km closer to the empirical resighting locations compared to the null model (range: 55–6793 km; 95% percentile interval 225–2352 km).

**Figure 2.**
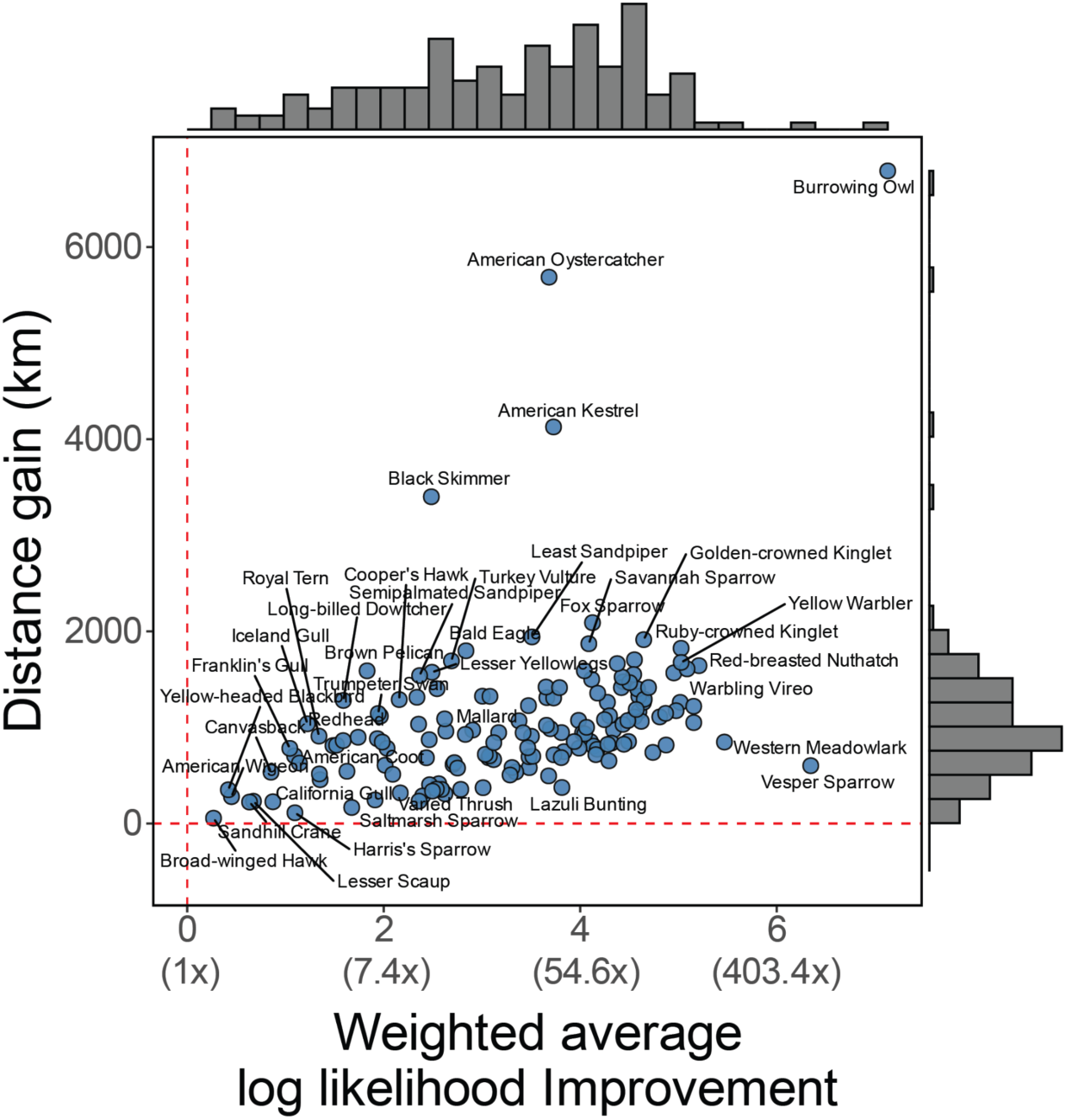
BirdFlow models for all 153 species perform better compared to a random sample from eBird Status & Trends (indicated by the dotted red line). Performance is evaluated by both weighted average log-likelihood improvement (x axis) and distance gain (y axis). Metrics are calculated across all hold-out transitions with different forecasting horizons. The log-likelihood improvement is additionally labeled using straightforward explanations (e.g., 7.4x means on average 7.4 times better in predicting the actual recovery location of the transitions compared to a random sample).

BirdFlow’s predictive performance decreased with increasing forecasting distance and time, but even at long temporal and spatial horizons, performance was better than a random sample from the S&T abundance surface (Fig. 3, a-b). Across time, most BirdFlow models continued to achieve better performance compared to S&T after 270 days (Fig. 3a) based on the relative distance gain metric. Across space, the averaged maximum functional distance beyond which the BirdFlow model was no longer favored over the null S&T abundance surfaces was 2449 km (95% percentile interval: 242–6395 km; Fig. 3 b-c). Similar results for the log-likelihood improvement metric can be found in Additional file 1: Fig. S6.

**Figure 3.**
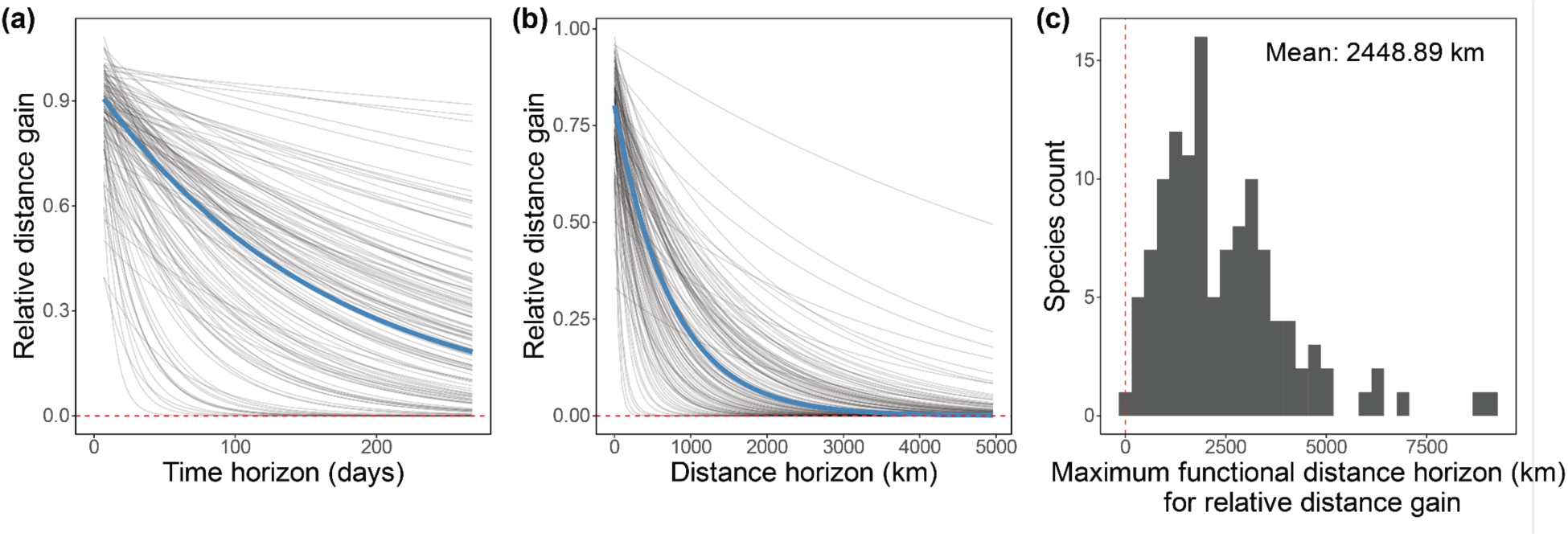
Model performance decays as time and distance horizon increase. Model performance decay is measured as the decrease of relative distance gain as the time horizon (a) and distance horizon (b) extends. Gray lines represent the fitted exponential regression function for each species, and blue lines represent the functions using cross-species-averaged parameters. (c) measures the maximum distance horizon that BirdFlow models still have advantages against a random sample from the abundance surface, defined as the horizon where the exponential decay curve (asymptotic to y=0) in (b) intersects with the y=0.05 line (5% closer to true resighting locations than S&T). Red dashed lines in (c) indicate zero distance horizon. See Additional file 1: Fig. S4 for schematic diagram of calculation in (c).

By comparing simulated trajectory statistics with empirical observations, we show that the biological characteristics of BirdFlow-generated routes are generally consistent with data from tracked individual birds (Fig. 4, Additional file 1: Fig. S7, Table S3). A biologically plausible model should expect around 95% of the empirically observed biological metrics to occur within the 95% simulated intervals and around 50% to occur within the 50% simulated intervals. For migration speed, on average 86% of the empirical observations fell within the 95% simulated intervals (range: 56%–100% across species), and 44% fell within the 50% simulated intervals (range: 22–58%) (Fig. 4). For route straightness, the averages are 90% (63%–100%) and 49% (11%–64%), respectively (Additional file 1: Fig. S7). For the number of stopovers, these values are 99.59% (97.14%–100%) and 98.44% (94.29%–100%), respectively (Additional file 1: Fig. S7). The results for the number of stopovers are less informative because the coarse weekly temporal resolution reduces the precision of stopover calculations.

**Figure 4.**
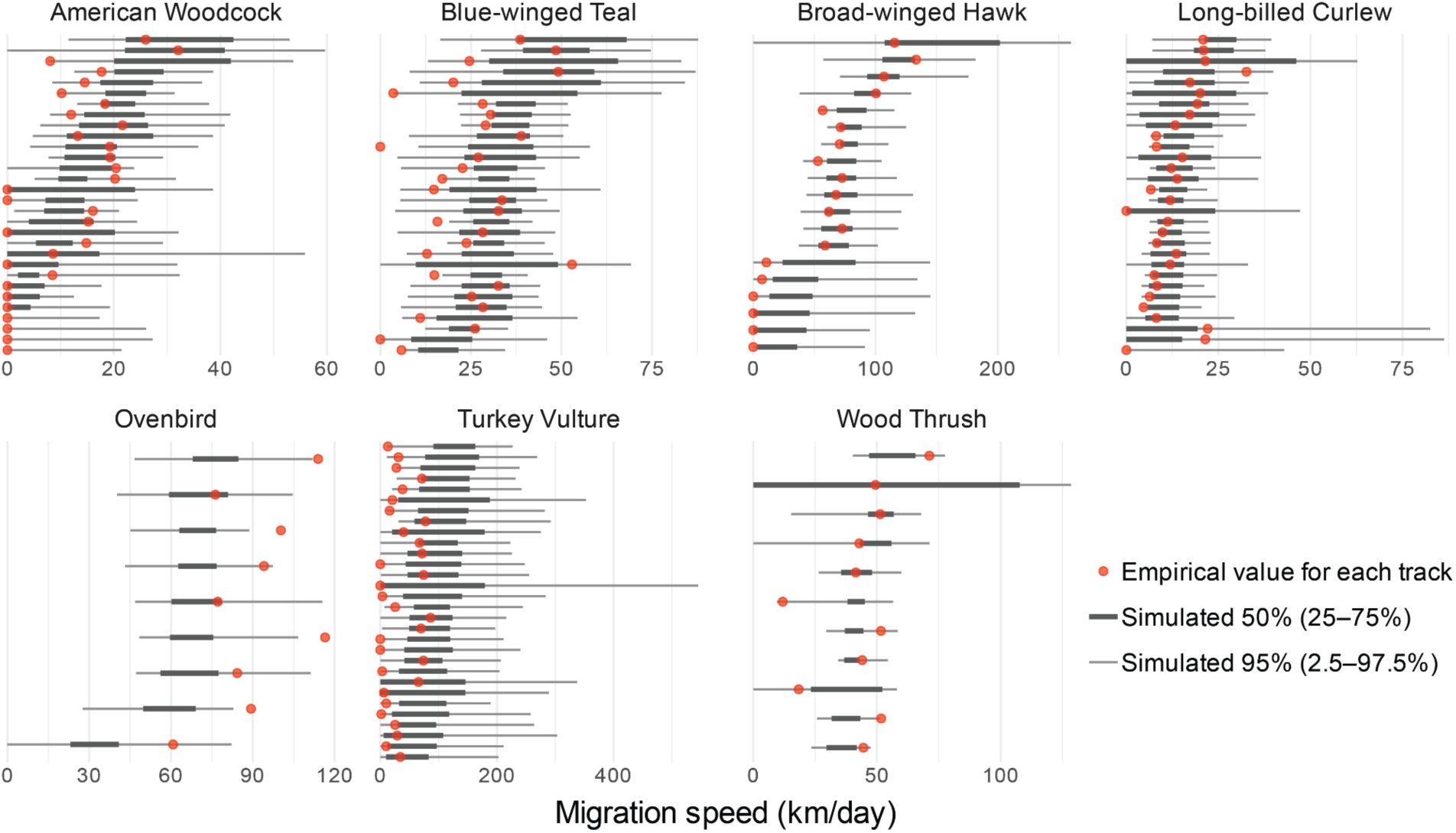
BirdFlow models simulate biologically plausible migration speeds. For each species, we randomly sampled at most 30 tracks from pre-breeding or post-breeding migration seasons. Red points indicate the empirical migration speeds calculated from GPS tracks and gray bars indicate 95% and 50% percentile intervals of the simulated migration speed. A quantitative summary of the figure can be found in Additional file 1: Table S3.

### Hyperparameter transfer among related species enhances model performance

The taxonomically informed model tuning methods showed better performance than the All-LOO method, both measured by relative distance gain and log-likelihood (Table 1). Models tuned using information from more closely related species performed better relative to the baseline All-LOO: Species-specific tuning performed the best (89.5% species showed log-likelihood improvement; 82.9% species show positive distance gain improvement), followed by Family-LOO (37.3% species showed log-likelihood improvement; 45.5% species show positive distance gain improvement) and Order-LOO (20.4% species showed log-likelihood improvement; 18.4% species show positive distance gain improvement).

**Table 1.**
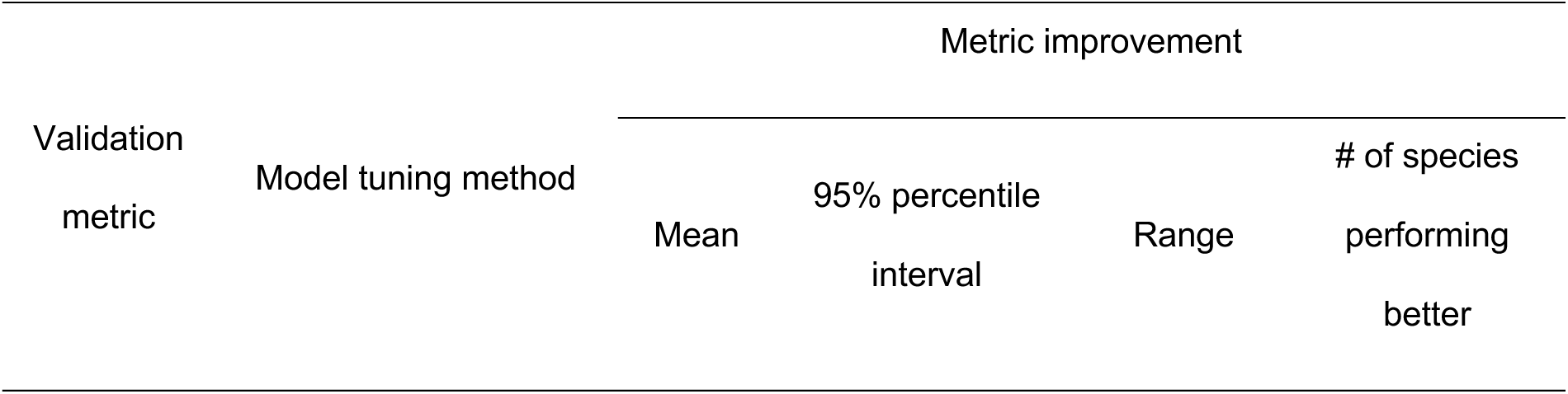
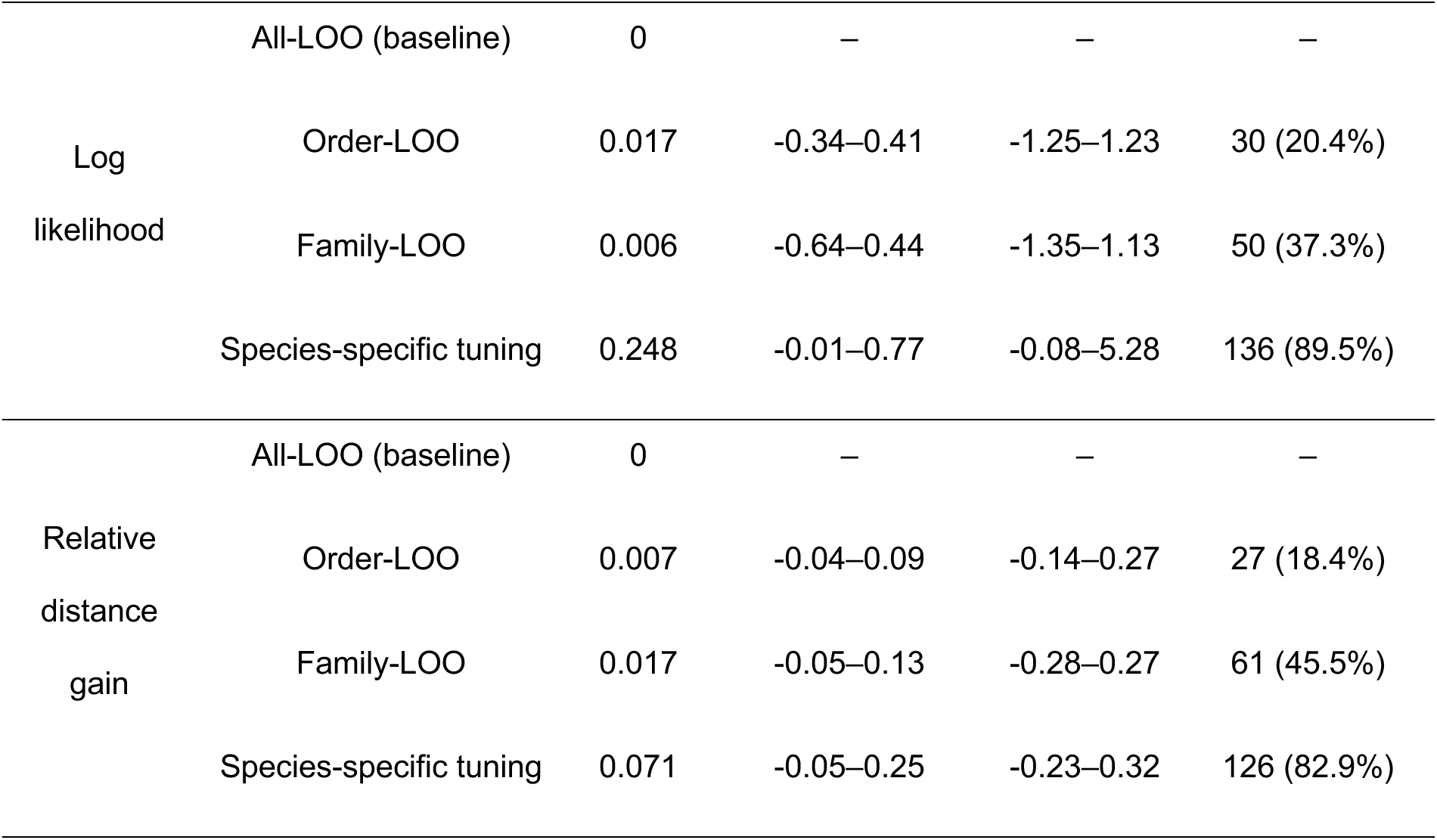
Taxonomically informed model tuning shows improvements over the all-leave-one-out method (ALL-LOO)

We further investigated the influence of transition data abundance on model tuning. The result shows that a small amount of training data during model tuning is sufficient for achieving a performance improvement compared to All-LOO baseline, and that more transition data for model tuning is not correlated with higher performance improvement across species (Additional file 1: Figure S8; Pearson’s r: 0.0012, Spearman’s r: 0.0942).

### Species traits explain variation in hyperparameters

While we did not observe statistically meaningful clustering patterns of hyperparameters by taxonomic group (Fig. 5), the regression results show that species’ morphological traits and distribution range summaries are predictive of their tuned hyperparameters (Additional file 1: Fig. S9a-c). A Random Forest regression model trained by 5-fold cross validation had moderate predictive power for the tuned values of hyperparameter 𝛽 (𝑅^G^ = 0.27), 𝜖 (𝑅^G^ = 0.20), and 𝛼 (𝑅^G^ = 0.27) across species. Body mass was the most important predictor of entropy weight and distance weight, with higher body mass correlated with higher entropy weights (Pearson’s r = 55; Additional file 1: Fig. S9d) and lower distance weights (Pearson’s r = -0.33; Additional file 1: Fig. S9f). Higher spatial variation of abundance and narrower range at breeding season were also correlated with higher distance weights (Pearson’s r = 0.45 and -0.47, respectively; Additional file 1: Fig. S9e, Fig. S5).

**Figure 5.**
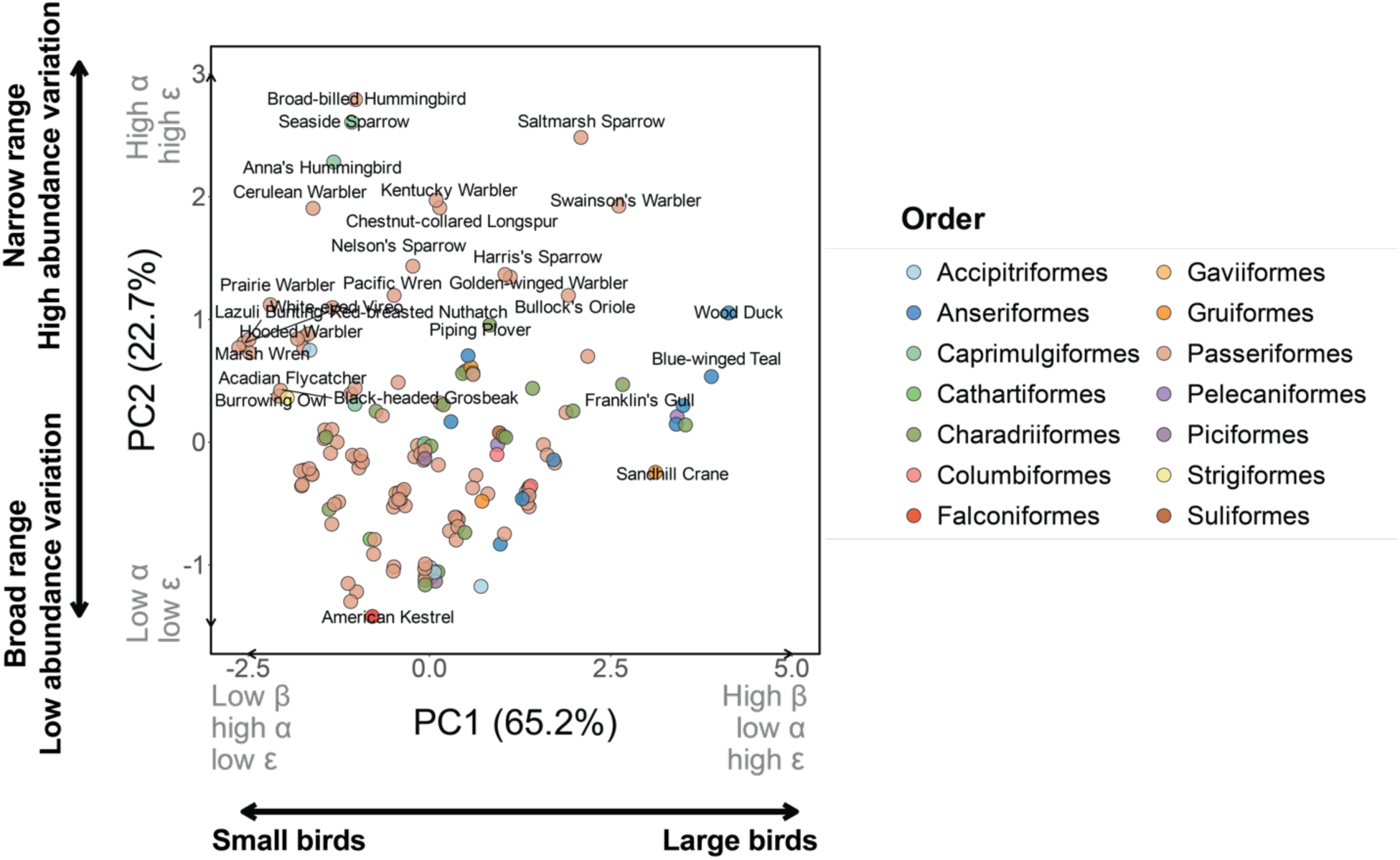
Associations between selected hyperparameters and species traits. Principal component analysis results for the hyperparameter patterns across species. Color denotes the order of the species. Gray text indicates the relative PC loadings of entropy weight 𝛽, distance weight 𝛼, and distance exponent 𝜖, respectively. Higher values in PC1 are correlated with higher entropy weight, lower distance weight, and higher distance exponent. Higher values in PC2 are correlated with higher distance weight and higher distance exponent. Small jitters in x and y axes were added for better visualization, since the best models for certain species share the same hyperparameter combination.

## Discussion

Our results show that BirdFlow models tuned with individual-based movement data can simulate routes that closely match actual individual tracks. By integrating the movements of a small sample of tracked individuals with eBird distribution data, we inferred range-wide population-level movement patterns for 153 species. These models provide a first quantitative yardstick for delineating population-specific migration routes, even when individuals are tracked or banded in only part of that range. Specifically, BirdFlow predicted movement transitions from week *i* to week *j* outperformed a null baseline of randomly sampling from a species’ abundance distributions corresponding to the resighting week. This result holds for >100 species (Fig. 2) across forecast horizons of a few thousand kilometers (on average 2449 km) and over 270 days (Fig. 3).

By integrating individual-based movement data from multiple sources into the hyperparameter tuning of BirdFlow, we greatly expand the number of species for which species-specific models can be tuned. Banding data alone enable tuning for 88.2% of the study species (defined as at least 20 transitions per species), but individual-level movement data remain scarce for many small-bodied passerines. Because of their low body mass, these species are difficult to track with current GPS technology [15, 25], and band-recovery data accumulate slowly due to low recovery rates [26]. In contrast, the Motus Wildlife Tracking System supports tuning for 49.7% of the study species, a smaller but rapidly growing fraction that is particularly valuable for small passerines [17]. As a result of their complementarity, combining banding and Motus data makes it possible to tune models for 98.7% of the studied species.

Our results demonstrate that improvements in model performance can be achieved with limited mark-resight data. Notably, simply increasing the number of individual transitions does not consistently lead to higher predictive performance across species (Additional file 1: Fig. S8).

This pattern is encouraging because it indicates that a small amount of empirical data can meaningfully calibrate and improve BirdFlow predictions for a focal species. This result also suggests it may be possible to increase the flexibility (i.e. number of hyperparameters) of the current BirdFlow model structure to learn more detailed patterns from the individual data, which could lead to increasing performance with higher amounts of tuning data.

To our knowledge, this study provides the first large-scale integration of distribution and individual-level data to infer animal movement across continents and among hundreds of species. As part of this study, we release the batch of BirdFlow models for 153 North American migratory species (see data statement), representing a comprehensive collection of population-level movement models spanning the full spatial and temporal ranges of these species.

Previous research on bird migration has developed quantitative and mechanistic models of movement and connectivity, but such approaches have typically been applied at the species level or to a limited number of intensively tracked populations. In contrast, BirdFlow resolves range-wide migration patterns at the population level across hundreds of species, addressing a key scaling gap given growing evidence that population-level variation in migratory strategies and population trends can be comparable to, or exceed, differences among species [10, 27, 28].

As expected, we see a decay of model performance along spatial and temporal forecasting horizons. A main explanation for this performance decay is the biological reality of populations mixing across space and time. For example, for species with low migratory connectivity, we expect the spatial connection patterns among populations to dissipate within a certain horizon [29]. For these species, BirdFlow models are not expected to be able to distinguish the movement of individuals in the same spatiotemporal unit. Alternatively, the decayed model performance could stem from the accumulation of prediction errors. Disentangling these two different factors requires further investigation into the migration connectivity estimation of the modeled species.

While most species’ models recovered the population-level characteristics of tracked individuals, some models exhibited consistent deviations. For example, the Ovenbird model consistently predicted lower migration speed and higher route straightness relative to empirical data, while the Turkey Vulture model consistently predicted higher migration speed and lower route straightness (Fig. 4, Additional file 1: Fig. S7). The biases indicate potential space to improve the parameterization and/or structure for those models.

Collectively, our results indicate that the three hyperparameters reflect species’ functional traits and distributional properties. For example, small passerine birds generally showed low entropy weight, high distance weight, and low distance exponent (Fig. 5; lower PC1 values; less spread and higher movement cost), while large birds in Anseriformes (ducks and geese) showed the opposite (higher PC1 values; more spread and lower movement cost). This result aligns with our understanding that small songbirds often depend on frequent flights and refueling, whereas large waterfowl accumulate substantial reserves and undertake long-distance flights [30–32].

We also find that species with narrow ranges (low spatial coverage) and high variation in spatial abundance show higher distance weight and higher distance power weight (higher PC2 values), resulting in a migration pattern with higher movement cost and favoring long jumps over small hops. On the other hand, species with broad ranges and low abundance variation show the opposite. This may occur because narrow-ranged species, being restricted to small geographic areas or specialized habitats, may face stronger migration barriers and have fewer suitable stopover habitats to choose from, which elevates the distance cost and forces long jumps rather than small hops (e.g., [33]).

The evidence of hyperparameter similarity among taxonomic groups opens the door for tuning species lacking any individual-level movement data. Hyperparameters could be transferred from close taxonomic relatives using a taxonomically informed leave-one-out approach, providing a practical fallback when species-specific movement data are unavailable. We show that BirdFlow models using hyperparameters transferred from phylogenetically adjacent species groups performed better than models using parameters transferred from distant or unrelated groups. In BirdFlow models, the three hyperparameters 𝛽, 𝛼 and 𝜖 encode different aspects of the migration pattern and strategies: the diversity of the migration routes, the cost of energy expenditure, and the trade-off between long-distance jumps versus short-distance hops. These diverse migration patterns and strategies are driven by species-specific trade-offs that ideally maximize the survival and reproduction of the species [34, 35]. Species that are closer in taxonomic relationship or functional traits might be more similar in migration patterns and strategies due to ancestral traits inheritance or convergent evolution [36] and consequently may also be closer in BirdFlow hyperparameter space. Despite this evidence, we did not find statistically significant clustering patterns of hyperparameters across orders or families, which indicates that within-group hyperparameter variation is comparable to among-group variation, further emphasizing that species-specific tuning is still important when available.

Additionally, because the validation framework is insensitive to both model structure and data type, it opens opportunities for additional refinement of the BirdFlow modeling framework, from adjusting hyperparameters to exploring alternative model formulations. For example, we acknowledge that log-likelihood may not be the optimal metric for hyperparameter tuning in all cases. Future work could therefore explore additional tuning criteria, such as distance gain or connectivity-based metrics. This tuning framework also provides a way to quantitatively evaluate different structures as more data become available, ensuring that results grow increasingly robust over time.

The probabilistic Markovian structure of BirdFlow models does have limitations in estimating the routes of individuals, in particular across long time scales and long distances. BirdFlow intrinsically models the synthetic movements of all individuals presented within a spatiotemporal unit, making it a model of space-time tracks along which the population’s biomass redistribution occurs. The Markovian nature of the current BirdFlow model algorithm precludes “memory” for individual movement histories. Accordingly, BirdFlow-simulated tracks are not expected to resolve all individual-level variation in the observed tracks. Rather, they represent the average movement patterns across all individuals at a given location and time.

While the generalization of movement patterns to species full populations and ranges is one of the key contributions of BirdFlow model output, we should be aware that the individual transition data on which models are tuned still have spatiotemporal biases. Tracking data often concentrate in a few locations rather than spanning the full species distribution [15], and banding or Motus records are largely restricted to sites where stations are established.

Moreover, relatively few transition data extend into Central and South America, which may bias the model to prioritize movement tuning for North America. While overfitting is unlikely with the current three-hyperparameter model, and models are heavily constrained by the fine-scale eBird distributional data, such biases warrant attention. If this tuning framework were applied to a more complex model structure, bias-control methods or balanced sampling strategies could be applied to quantify such potential bias.

Another limitation of this work, and of BirdFlow more generally, is that it does not fully propagate uncertainty in eBird S&T relative abundance surfaces and tracking observation inputs (e.g. cf. Buderman et al. [37]). Rather than jointly estimating movement processes and observation models from multiple data streams within a single hierarchical model, BirdFlow uses abundance surfaces as a ground truth, while additional data sources such as tracking and mark-resight data are used primarily to inform model tuning and evaluation. As a result, uncertainty is not explicitly propagated throughout the full inference process. In this study, we did not incorporate uncertainty in the eBird S&T abundance estimates because doing so at this scale would have been computationally prohibitive. Although the criterion requiring mean correlation score > 0.98 may provide some tolerance to uncertainty and help reduce overfitting during model tuning, it does not substitute for formal uncertainty propagation. Future extensions could more formally account not only for uncertainty in the S&T abundance surfaces, but also for observation-process biases, such as spatiotemporal bias in banding and Motus data. Such approaches may provide stronger uncertainty quantification and a clearer representation of heterogeneity among individuals or populations.

As illustrated in Fig. 6, species-specifically tuned BirdFlow models provide an approach for addressing a broad range of ecological, evolutionary, and conservation questions. These models provide a population-resolved, continent-scale representation of migratory movement across the annual cycle, making it possible to link movement dynamics to environmental context and to quantify population differences without collapsing them into a single species-level pattern. This population-level resolution also allows direct, comparable tests of how migration strategies diverge or converge across populations and taxa. Finally, the same framework supports conservation-relevant attribution by connecting seasonal ranges, enabling researchers to evaluate which times of year and which regions are most strongly associated with among-population differences in trends and therefore most informative for explanation, prediction, and targeted intervention (Fig. 6). These questions often require range-wide coverage as provided by BirdFlow and are difficult to address with distributional data or sparse individual-tracking data alone. Together, these advances position BirdFlow as a general framework for integrating multi-source data to estimate range-wide bird movements at continental scales, opening new opportunities to link population ecology with applied management across hundreds of species.

**Figure 6.**
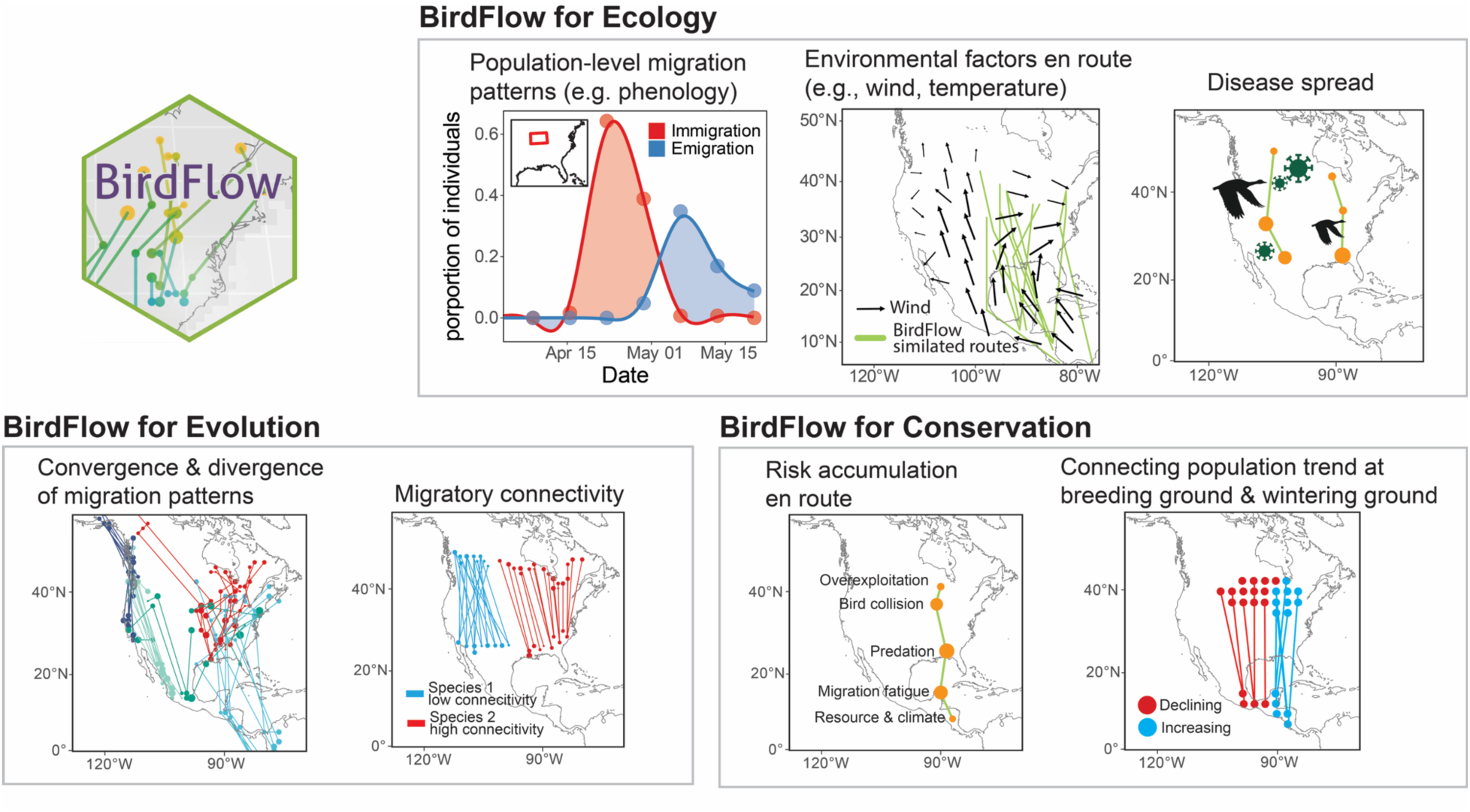
Applications of using BirdFlow for ecological, evolutionary, and conservation research. In ecological research, BirdFlow enables (i) quantification of population-level migration dynamics, such as regional influxes and effluxes, (ii) integration of migration patterns with environmental conditions to assess environmental drivers and barriers of movement, and (iii) linkage with surveillance data to study the spread, control, and prevention of wildlife-borne diseases. In evolutionary research, BirdFlow-simulated trajectories support analyses of (i) convergence and divergence in migration routes within and among species and (ii) variation in migratory connectivity, providing insights into the mechanisms shaping differences in migration strategies. In conservation applications, BirdFlow models facilitate (i) assessment of population-specific anthropogenic and environmental risks accumulated along migration routes and (ii) identification of the seasons and locations most influential for population trends by connecting breeding and wintering grounds.

## Conclusions

By integrating eBird Status & Trends abundance surfaces with multi-source mark–resight observations, including banding recoveries, Motus telemetry, and GPS tracking, we parametrized BirdFlow models for 153 North American migratory bird species. This continent-wide inference of population movements over the full annual cycle provides a quantitative basis for delineating population-specific routes across species’ full ranges, complementing existing individual-based tracking efforts and helping fill a key gap in ecological research and applied conservation. Tuned models consistently outperform abundance-based null expectations, remain informative across forecasting horizons of thousands of kilometers and many months, and generate biologically realistic migration routes that align with individual movement data. When species-specific movement data are unavailable, taxonomically informed hyperparameter transfer enables improved model, demonstrating that migration strategies share partial predictability across related taxa. Together, these advances establish BirdFlow as a general, extensible framework for estimating the comprehensive migratory movements of entire populations for hundreds of species, offering a foundation for more accurate predictions for ecological understanding, conservation planning, disease surveillance, aviation risk assessment, and public outreach as new data continue to accumulate.

## List of abbreviations

𝐿𝐿: log-likelihood
𝛥𝐿𝐿: weighted log-likelihood
eBird S&T: eBird Status & Trends relative abundance surfaces
LOO: leave-one-out
BF: BirdFlow
𝑝_12_: the probability assigned by the BirdFlow model to a possible destination in space and time, conditional on a specified starting location and time.

## Declarations

### Ethics approval and consent to participate

Not applicable

## Consent for publication

Not applicable

## Availability of data and materials

Code is available at https://github.com/birdflow-science/BirdFlowTuning. Data and the species-specifically tuned models are available at https://doi.org/10.5281/zenodo.17545007. Models can be loaded using the BirdFlowR package (https://github.com/birdflow-science/BirdFlowR). Tuned models that passed manual quality control will be released along with the BirdFlowR package in the future.

## Authors’ contributions

Conceptualization: DS, YC, DLS, BVD, AMD, EP, MF; Data curation: LEB, SAM, DLS, BVD, YC, EP, YD, AMD; Formal analysis and investigation: YC, DLS, EP, YD; Methodology: DS, YC, DLS, EP, BVD, AMD, MF, DF; Software: DLS, EP, YC; Validation: YC, DLS, EP, YD. Visualization: YC. Project administration and funding: DS, AMD, BVD; DF; Supervision: BVD, AMD, DS; Writing: All authors.

## Acknowledgements

We thank the USGS bird banding lab for the bird banding data collection and Birds Canada for preparing and sharing the data of the Motus Wildlife Tracking System and the individual contributors to these projects. We are also grateful to the many thousands of participants, reviewers, and partner organizations around the world who support and contribute to eBird.

## Supplementary Figures

**Figure S1.**
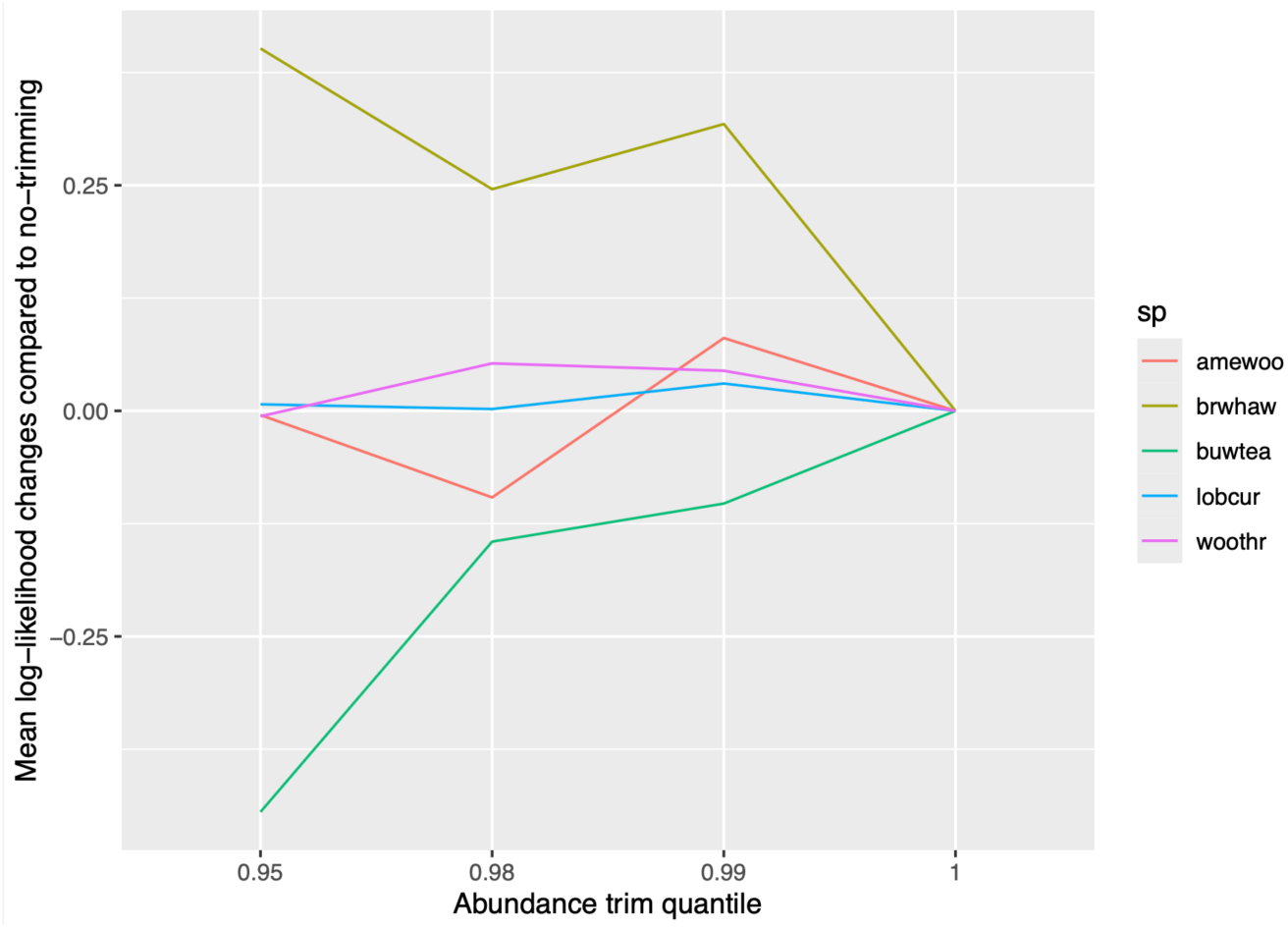
Sensitivity analysis of trimming quantile choices. To avoid the influence of high-abundance outliers, we truncated abundance distributions by replacing all values above the 0.99 quantile with the 0.99 quantile value. We showed that this outlier removal improved the average log-likelihood across the 225 models for 4 out of 5 species tested, for which individual tracking data were available at the time of analysis. The multi-species averaged log-likelihood improvement was the highest with 0.99 quantile trimming (𝛥𝐿𝐿 = 0.074), followed by 0.98 (𝛥𝐿𝐿 = 0.012), 1 (no trimming; 𝛥𝐿𝐿 = 0), and 0.95 (𝛥𝐿𝐿 = -0.009).

**Figure S2.**
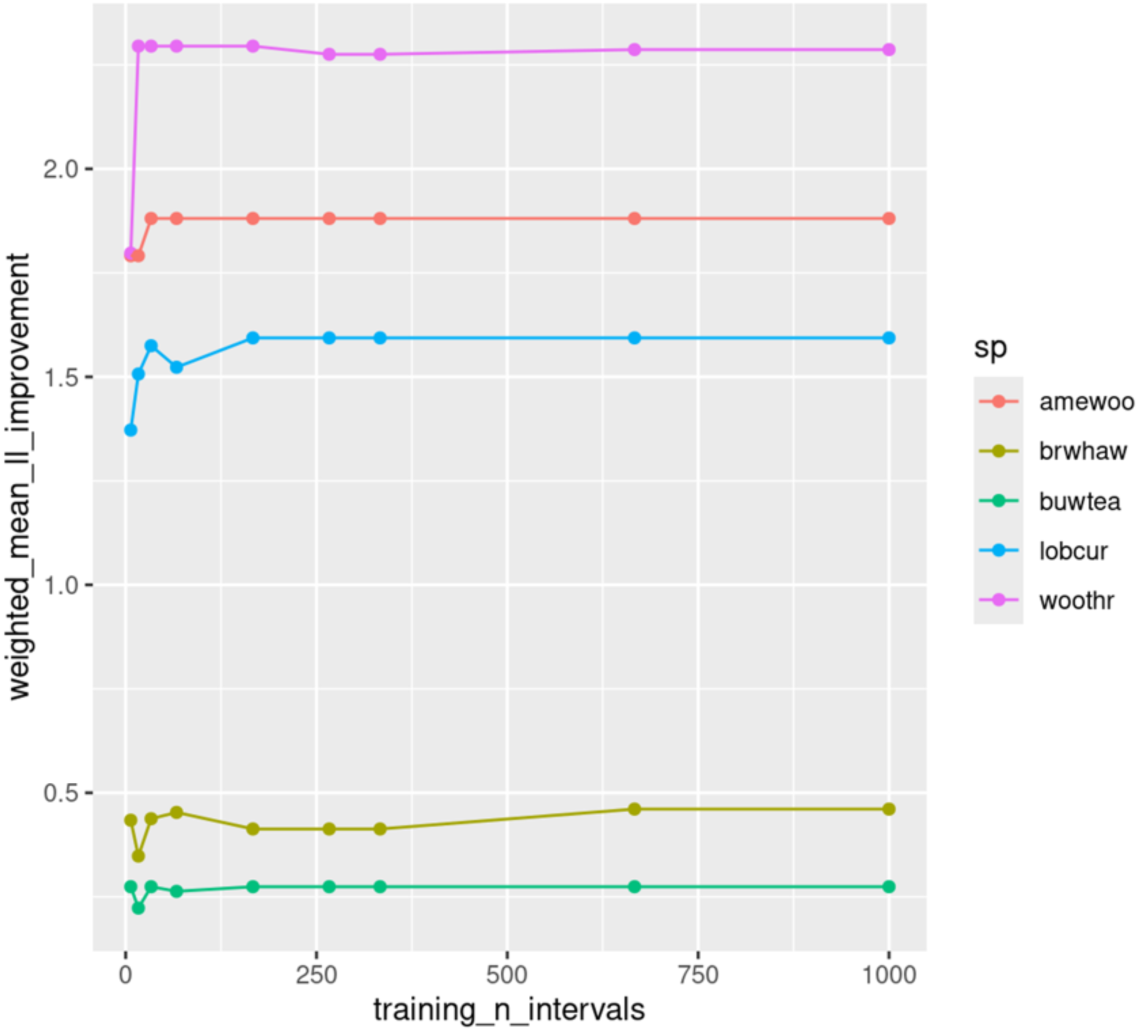
Sensitivity analysis of number of training transitions. To determine the minimum number of transitions required to train BirdFlow models, we applied a learning-curve analysis for five species for which individual tracking data were available at the time of analysis, each trained with increasing number of transitions. Model performance was evaluated on a hold-out test dataset as described in the Method section. We found that models trained with approximately 20–50 transitions achieved substantially improved performance (log-likelihood) and began to approach a performance plateau. We therefore chose a threshold of 20 transitions to train the models for maximum number of species.

**Figure S3.**
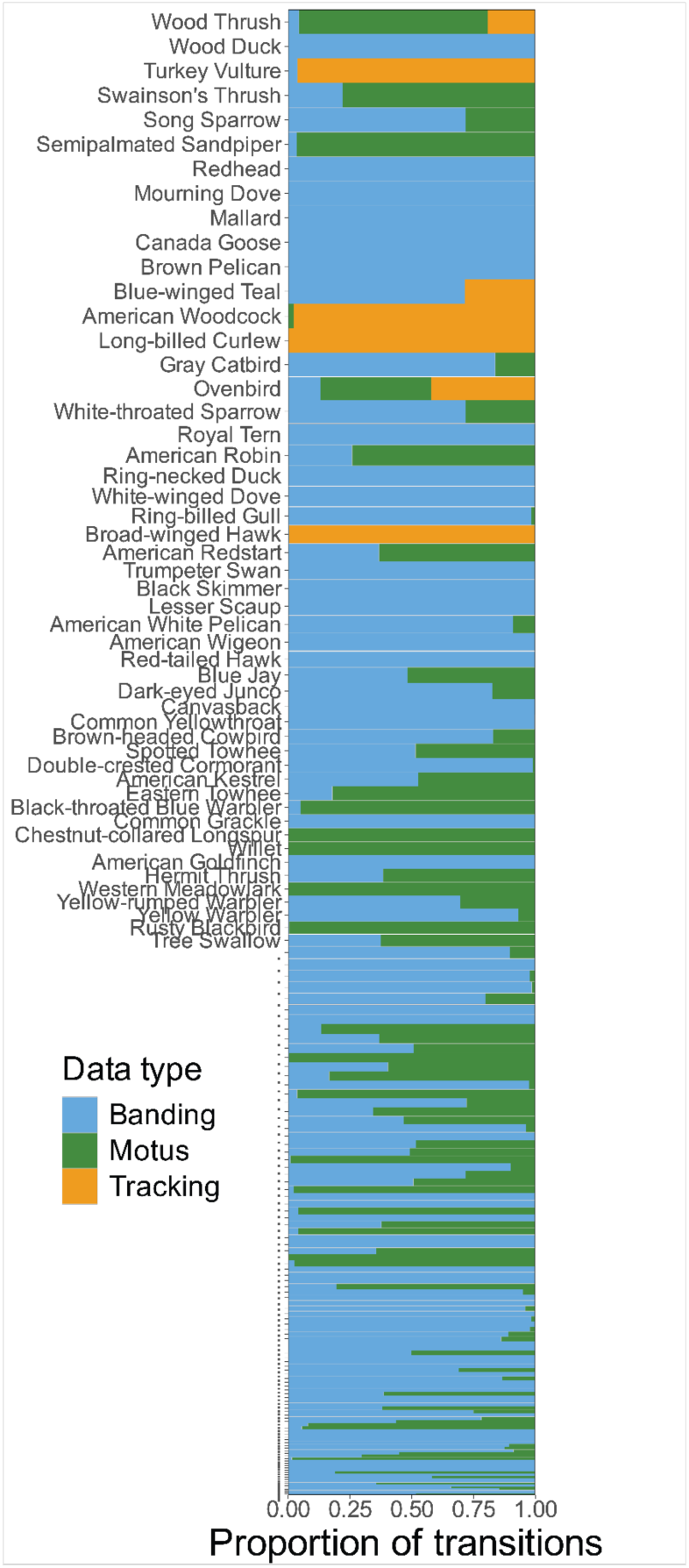
The transition data composition for each species, colored by data sources. The height of each block is proportional to the square root of total transition count available for the species. The width of the block represents the proportion of transitions from each data source for each species. The top 50 species with most abundant transition data are labeled with their eBird English common name.

**Figure S4.**
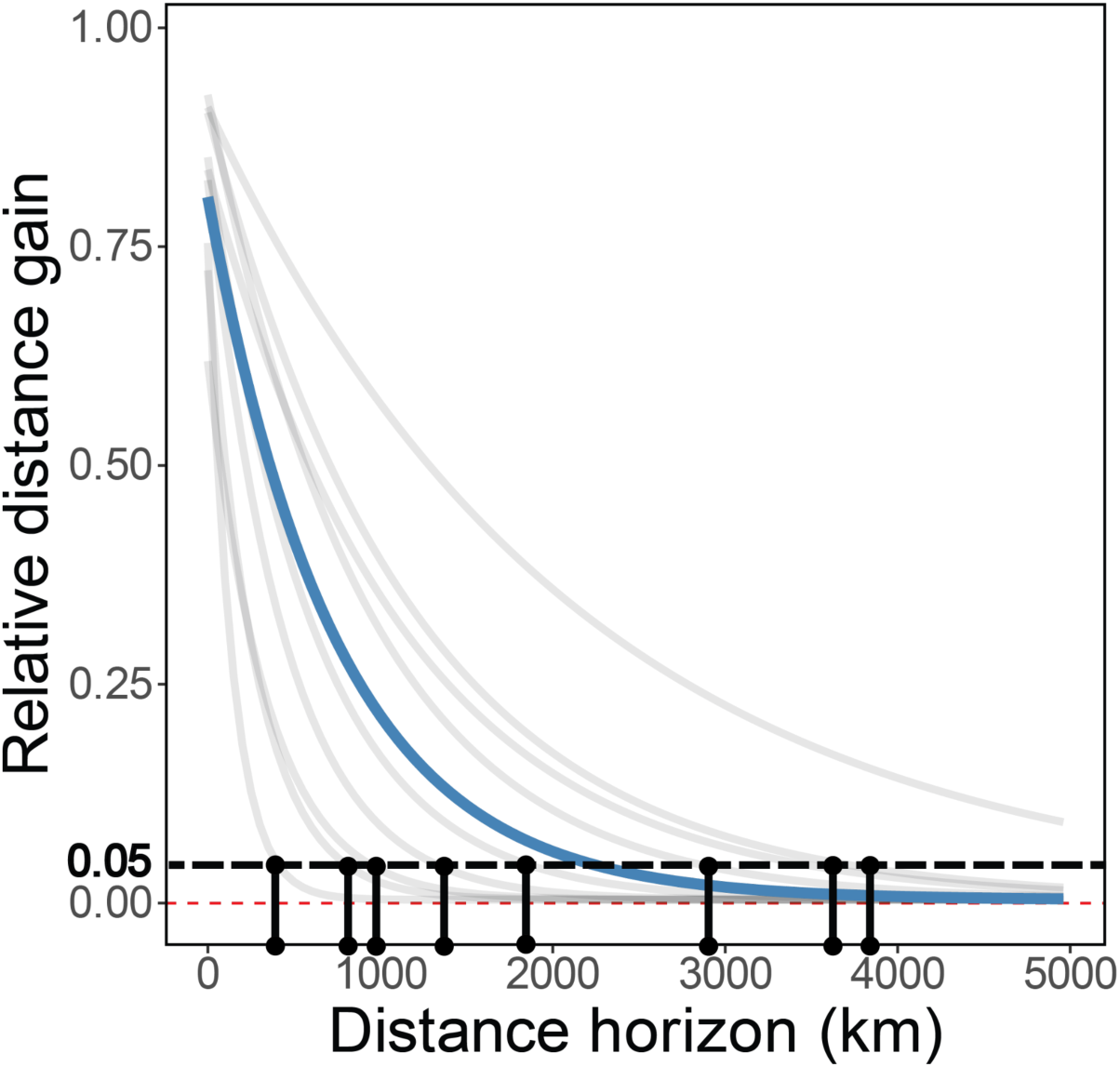
A schematic diagram of how the maximum functional forecasting horizon is calculated. We set a threshold of 0.05 for both relative distance gain and log-likelihood improvement, with values below the threshold being considered as BirdFlow models having no advantages over the baseline. We calculated the intersection of each species’ model performance decay line with the threshold, and averaged the distance horizon at the intersection as a multispecies averaged maximum functional horizon.

**Figure S5.**
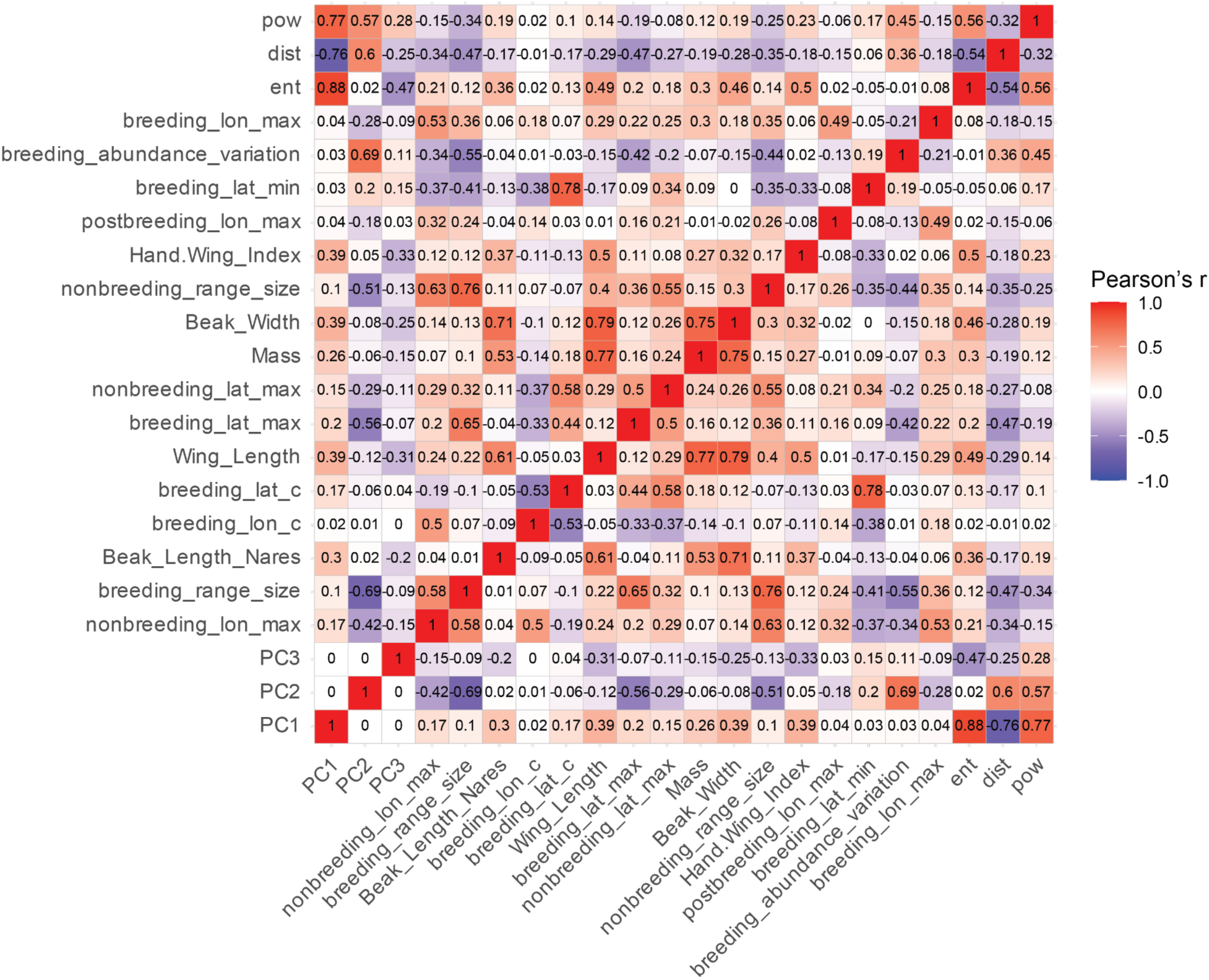
Pairwise Pearson’s correlation coefficients among variables included in the Random Forest regression. The variables include the hyperparameters, principal components of the hyperparameters, species range summaries, and species morphological traits. Each datapoint is a species. Species range summaries and morphological traits with Pearson’s correlation coefficients over 0.8 were considered highly collinear, and in such cases only one variable was retained while the other was removed before the analysis.

**Figure S6.**
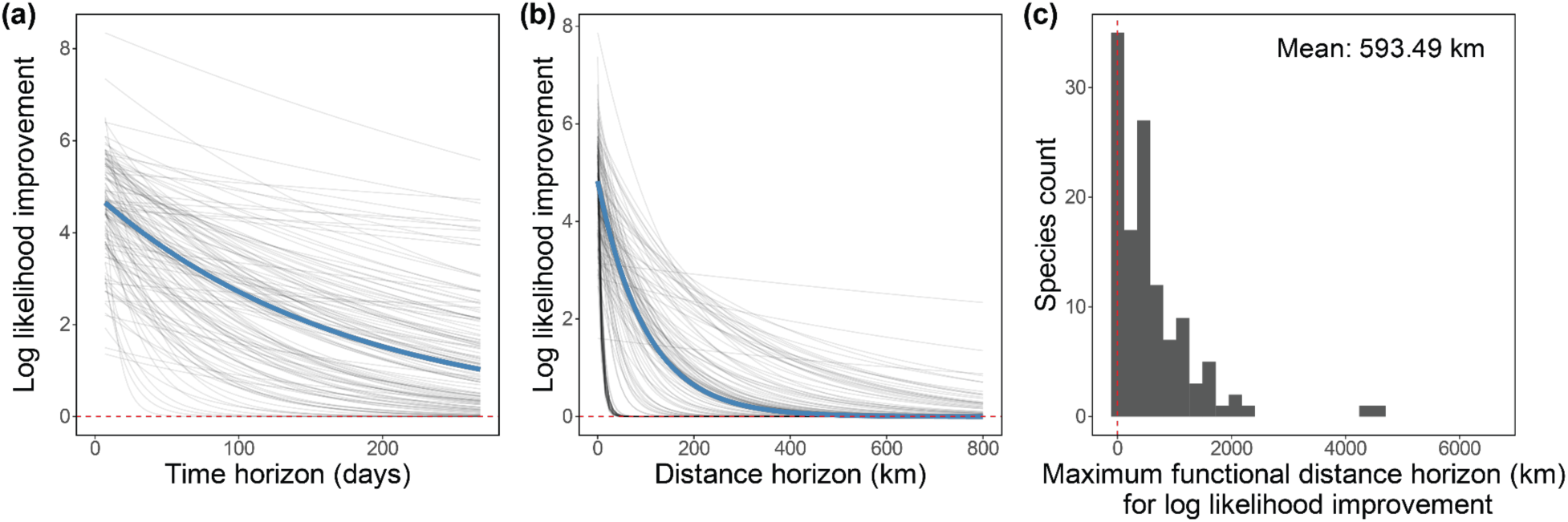
Model performance decays as time and distance horizon increase. Model performance decay is measured as the decrease of log-likelihood as the time horizon (a) and distance horizon (b) extends. Gray lines represent the fitted exponential regression function for each species, and blue lines represent the functions using cross-species-averaged parameters. (c) measures the maximum distance horizon that BirdFlow models still have advantages against a random sample from the abundance surface in terms of log-likelihood across species, defined as the horizon where the exponential decay curve in (b) intersects the y=0.05 horizontal line (1.05 times better than S&T). Red dashed lines in (c) indicate zero distance horizon. See Supplementary Fig. 4 for schematic diagram of calculation in (c). On average, the maximum functional distance horizon is 593.49 km for log-likelihood improvement (95% percentile interval: 31 km - 2145 km). The fact that the maximum functional distance horizon for log-likelihood improvement (593 km) is lower than that for relative distance gain (Fig. 3) demonstrates that log-likelihood is less tolerant to error accumulation along the prediction horizon, since it only measures the performance at the exact resighting location and does not consider geographic closeness.

**Figure S7.**
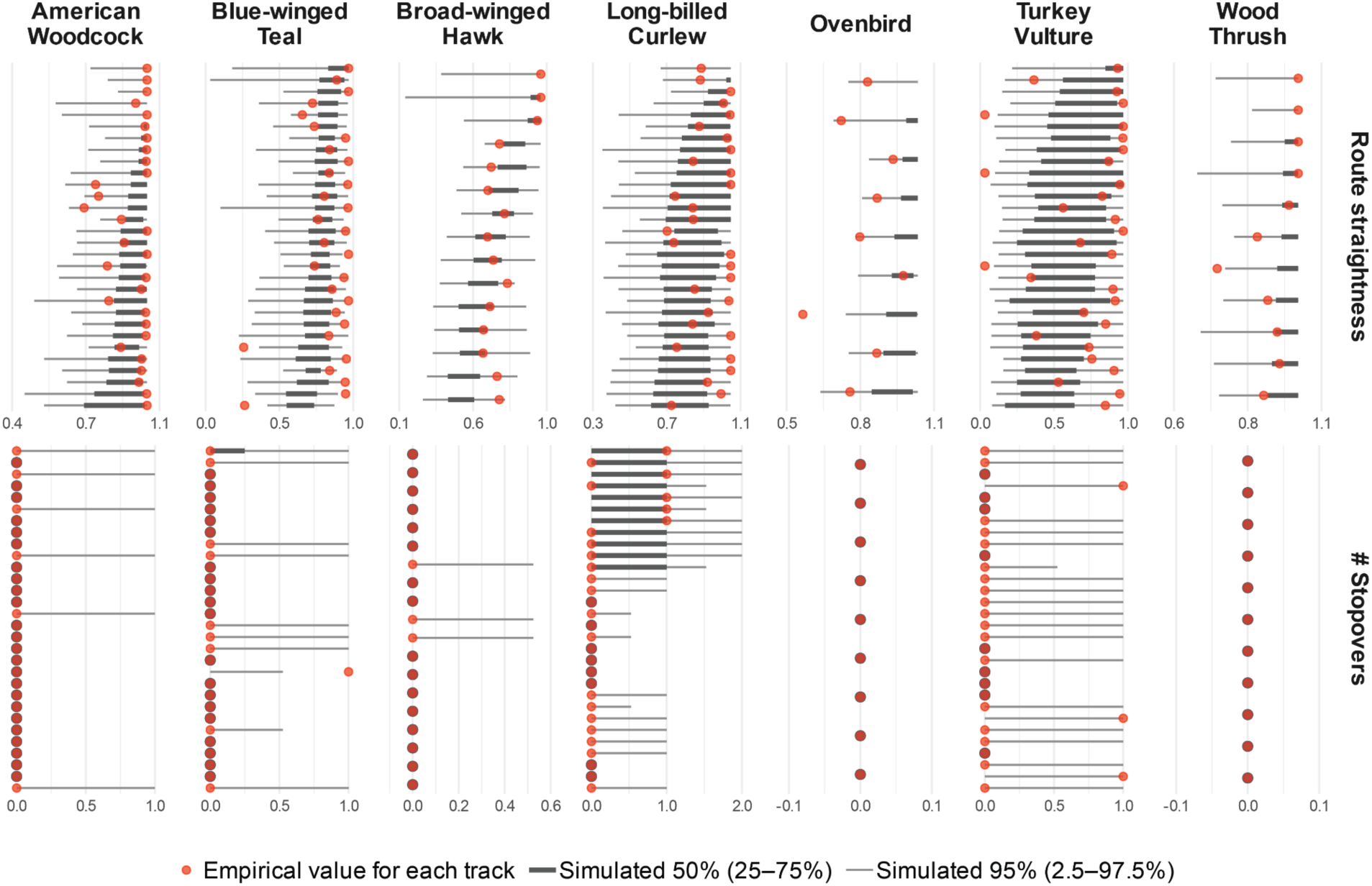
BirdFlow models simulate biologically plausible route straightness and number of stopovers. For each species, we randomly sampled at most 30 tracks from pre-breeding or post-breeding migration seasons. Red points indicate the empirical metrics calculated from the GPS tracks and the gray bars indicate 95% and 50% percentile intervals of the simulated metrics. A quantitative summary of the figure can be found in Additional file 1: Table S3.

**Figure S8.**
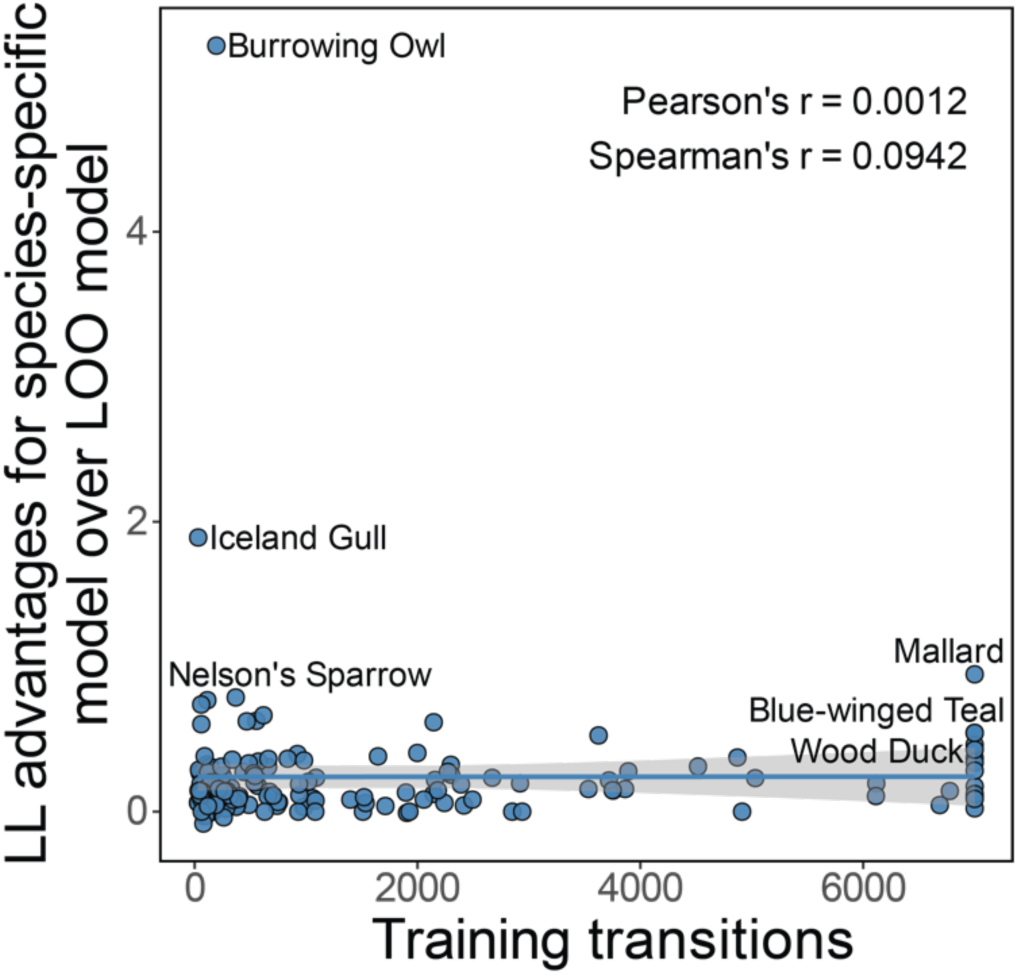
Lack of correlation between the improvements of species-specifically tuned models and the amount of training transitions. Although species-specific tuning improves model performance compared to All-LOO baselines, more training data does not necessarily lead to higher log-likelihood improvement across species. Each data point represents a best model for each species tuned on species-specific data.

**Figure 9.**
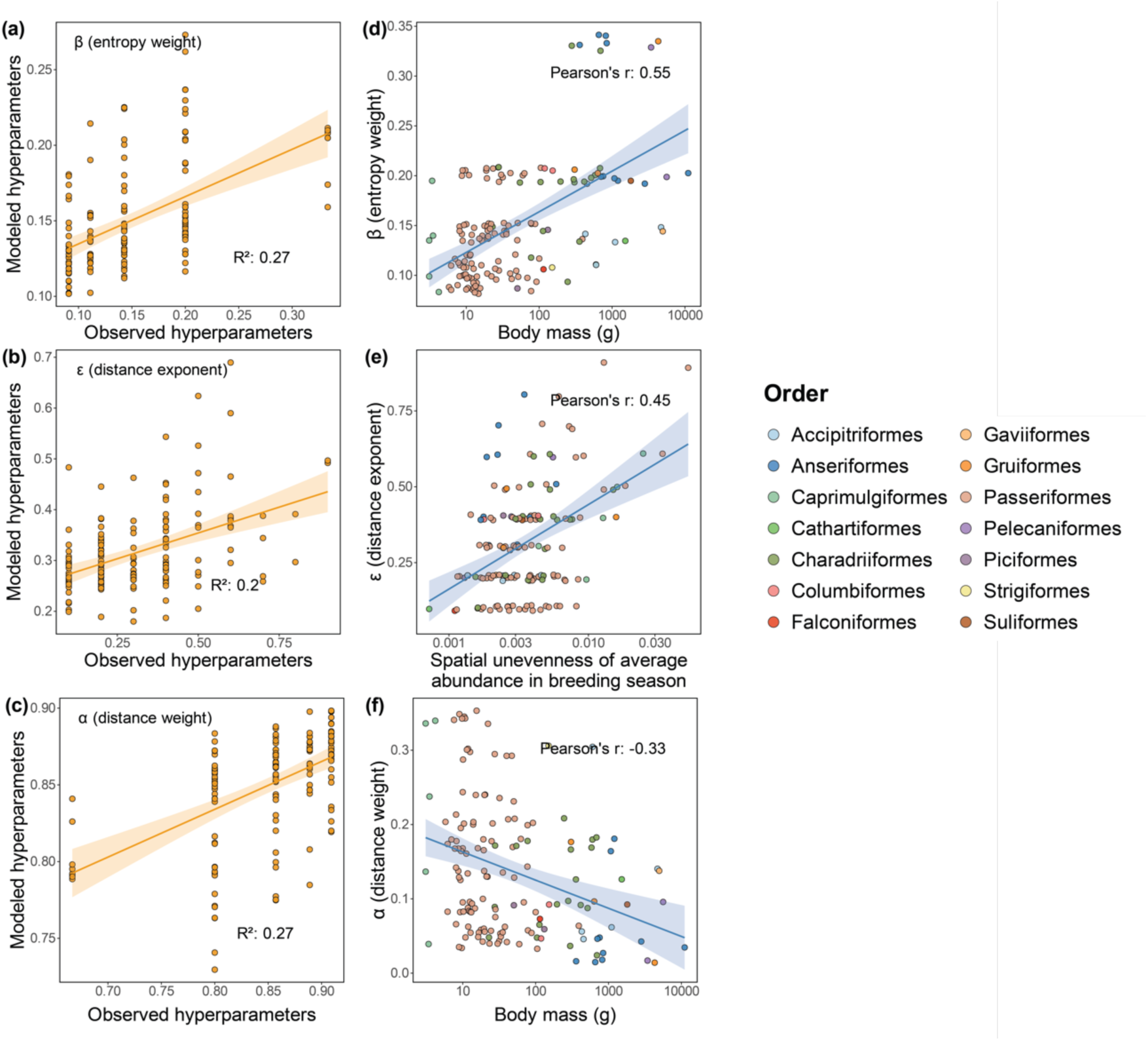
Explaining hyperparameters using species traits. We trained Random Forest models using species morphological traits and range summary statistics to explain the hyperparameter patterns among species. The Random Forest model was trained using 5-fold cross validation, and the out-of-fold prediction results for all species are shown for hyperparameter 𝛽 (a), 𝜖 (b) and 𝛼 (c), respectively. The hyperparameters are also linearly regressed against the explanatory variables with the highest feature importance score in the Random Forest models for 𝛽 (d), 𝜖 (e) and 𝛼 (f). The values on the horizontal axes are log-transformed to calculate the correlation coefficients in (d-f). Small jitters in x and y axes were added in (d-f) to disperse overlapping points.

**Figure S10.**
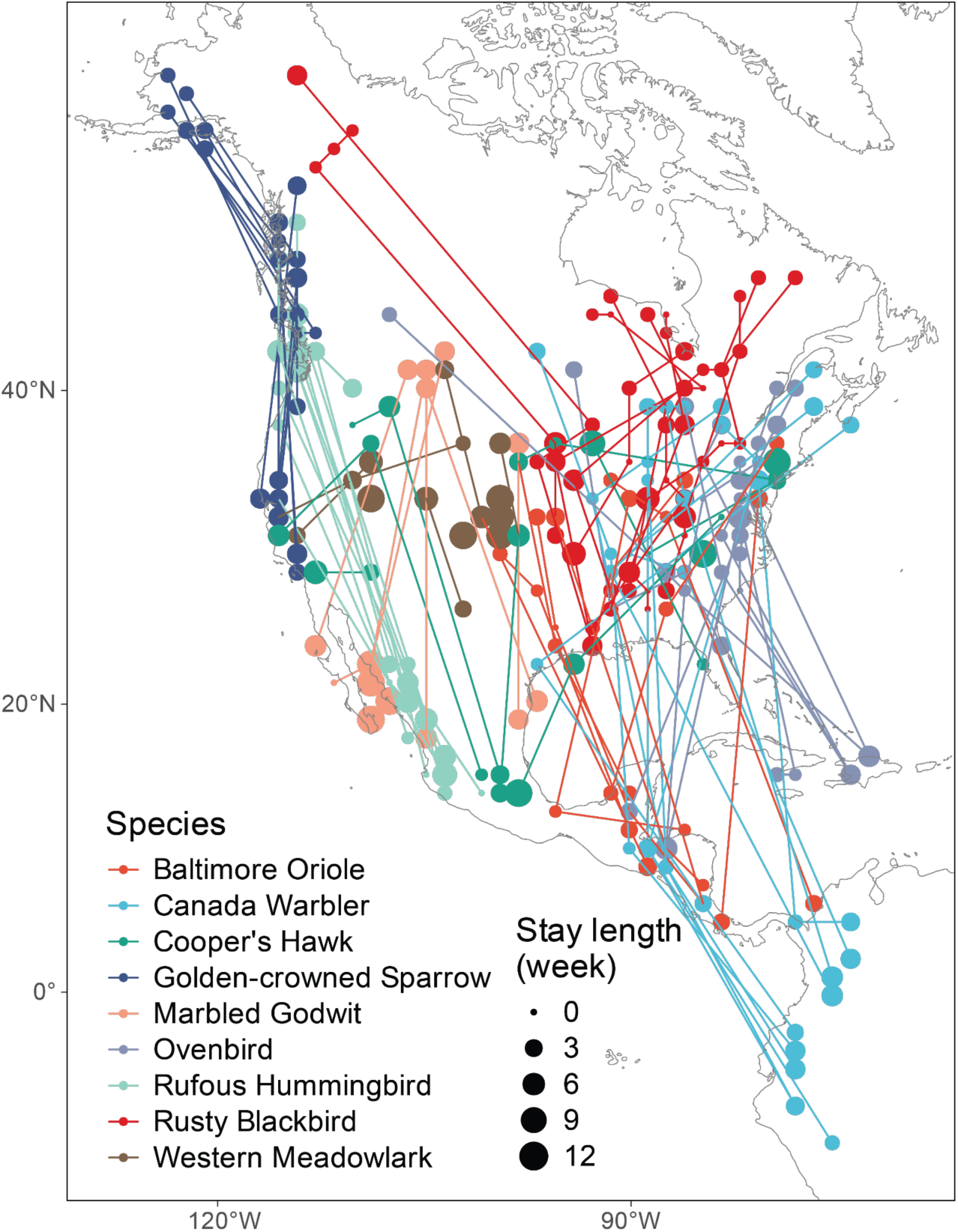
The BirdFlow-simulated spring migration routes for 9 migratory species. Stay length indicates the inferred total length of stay at the stopover sites in weekly temporal resolution, with zero stay length indicating non-stopping or stopover shorter than one week. 10 routes were simulated for each species.

## Supplementary Tables

**Table S1.** The sources of tracking data for each species used in this research.

| Common name | Scientific name | Data source |
| --- | --- | --- |
| American Woodcock | *Scolopax minor* | Movebank ID: 12170798, 351564596 |
| Turkey Vulture | *Cathartes aura* | Movebank ID: 16880941 |
| Ovenbird | *Seiurus aurocapilla* | Movebank ID: 205765709 |
| Long-billed Curlew | *Numenius americanus* | Movebank ID: 42451582 |
| Blue-winged Teal | *Spatula discors* | Movebank ID: 446107870 |
| Broad-winged Hawk | *Buteo platypterus* | Movebank ID: 28691134 |
| Wood Thrush | *Hylocichla mustelina* | Supplementary material of Stanley et al. 2015 |

**Table S2.**
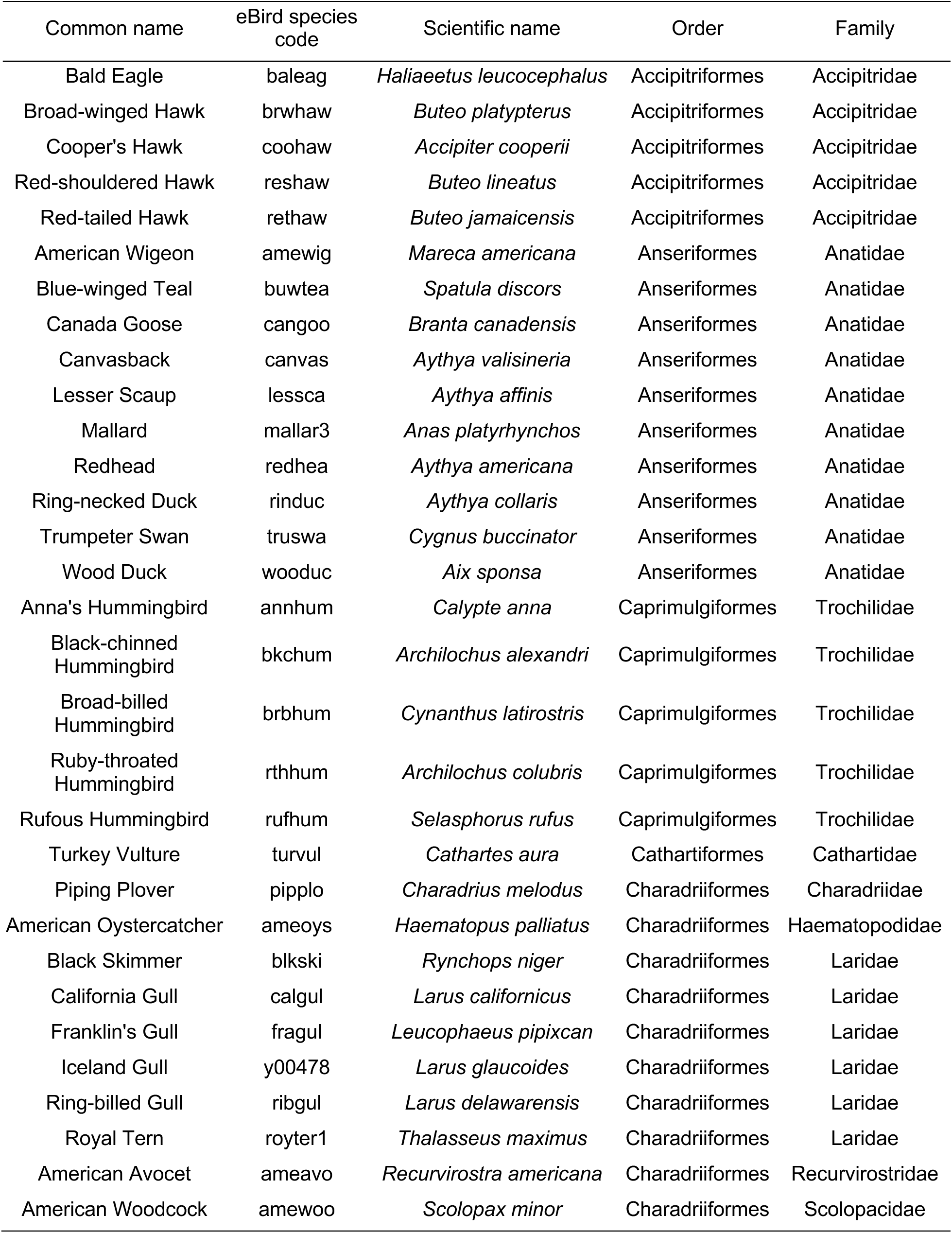
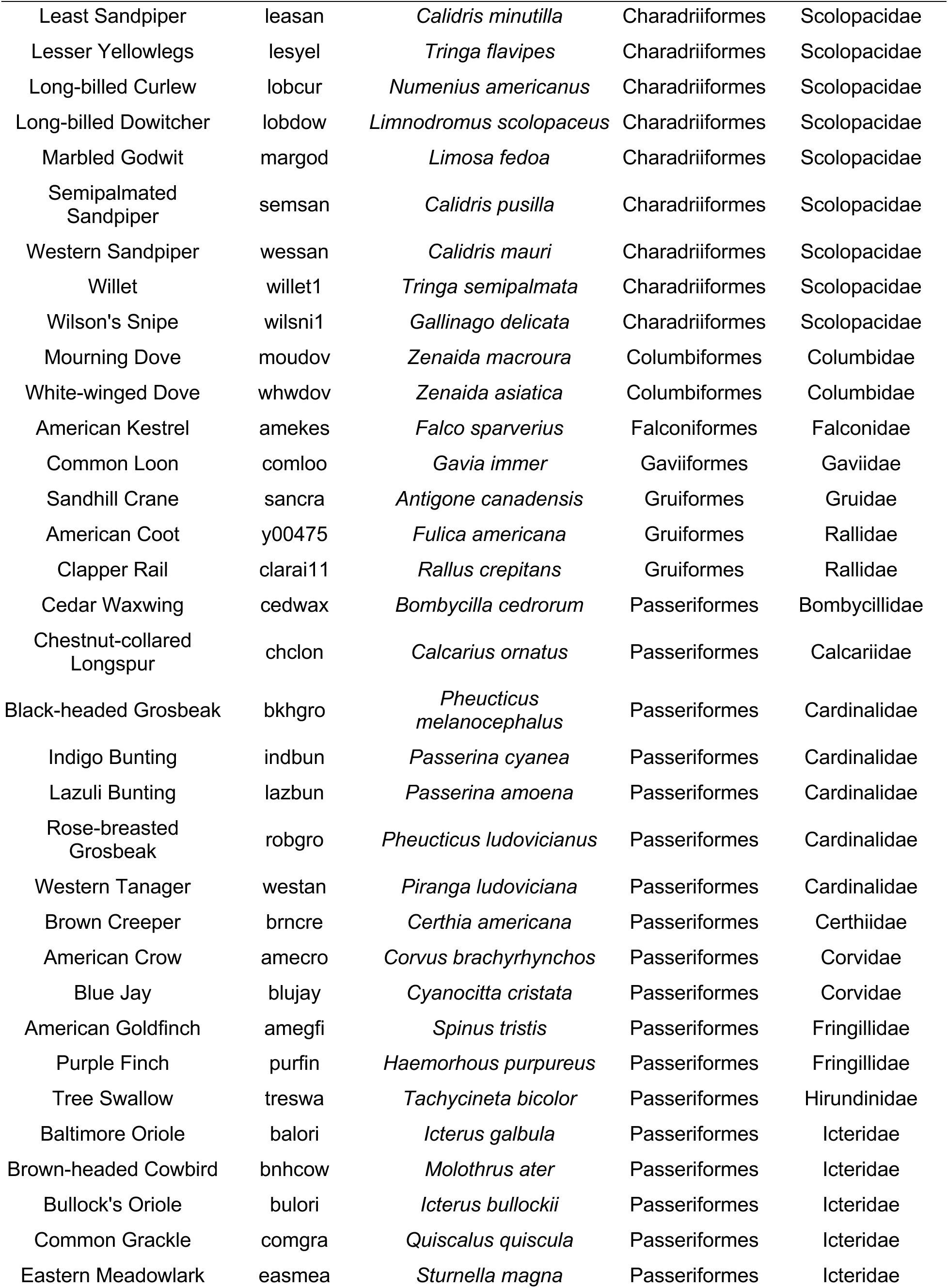
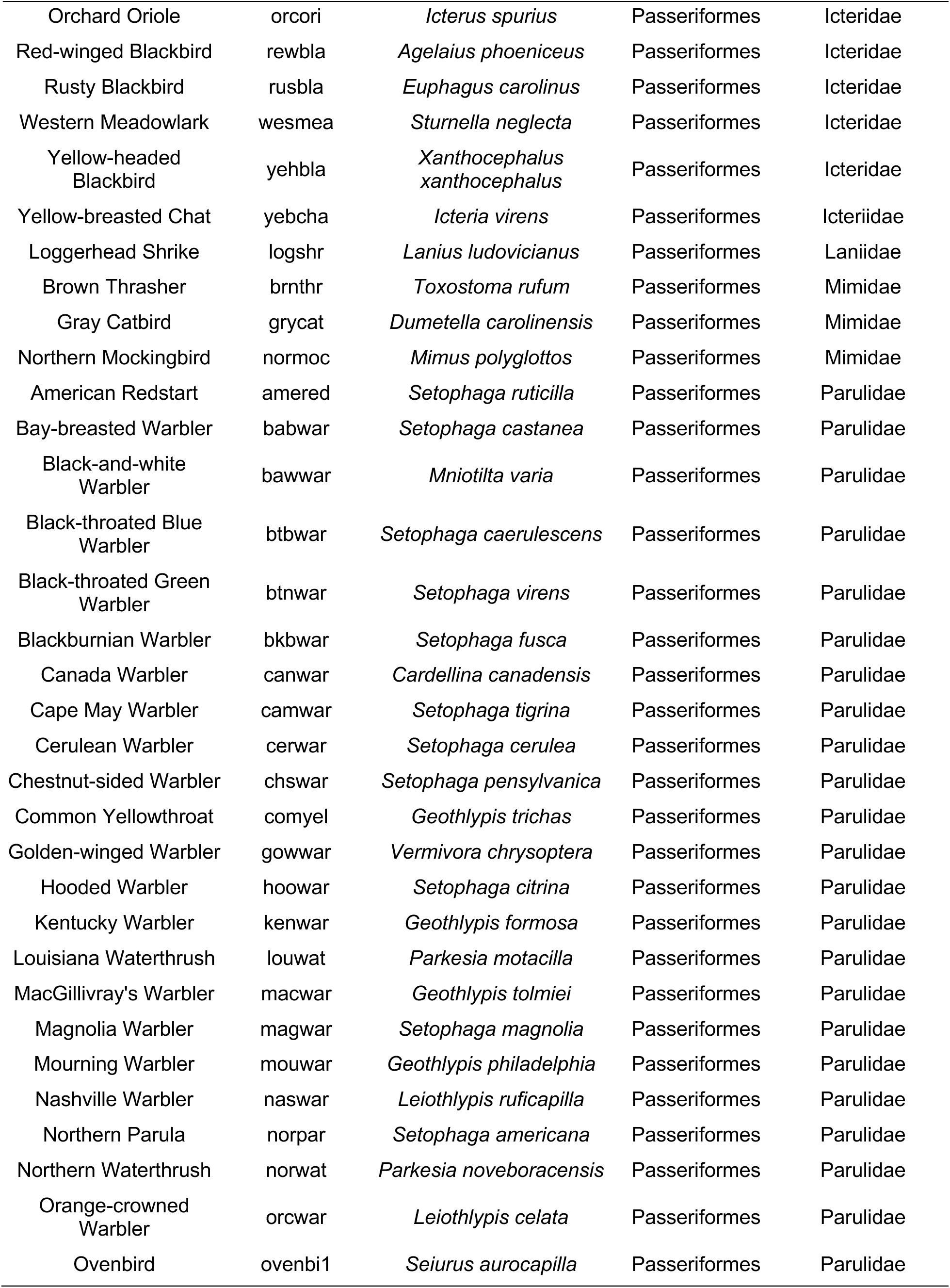
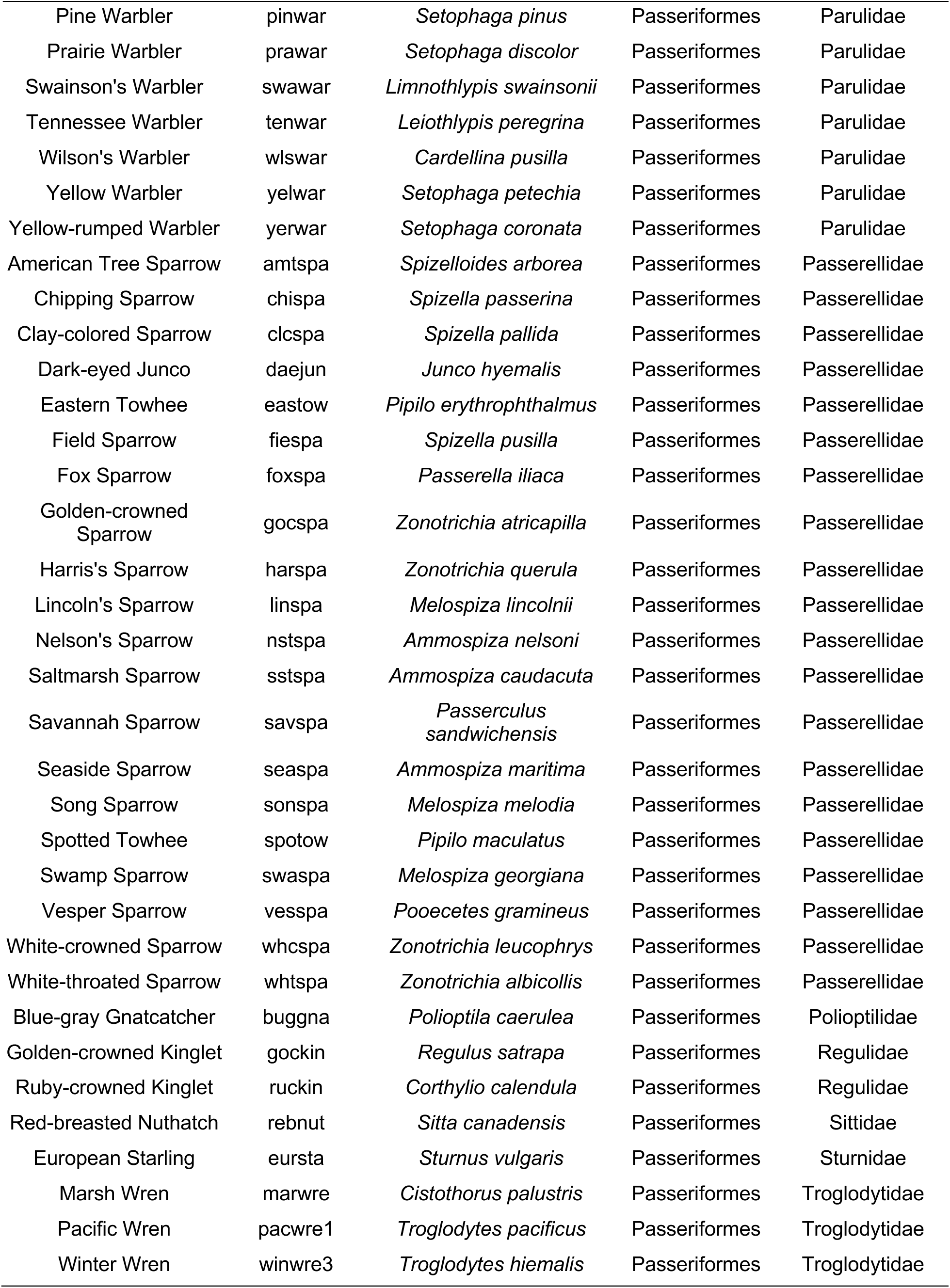
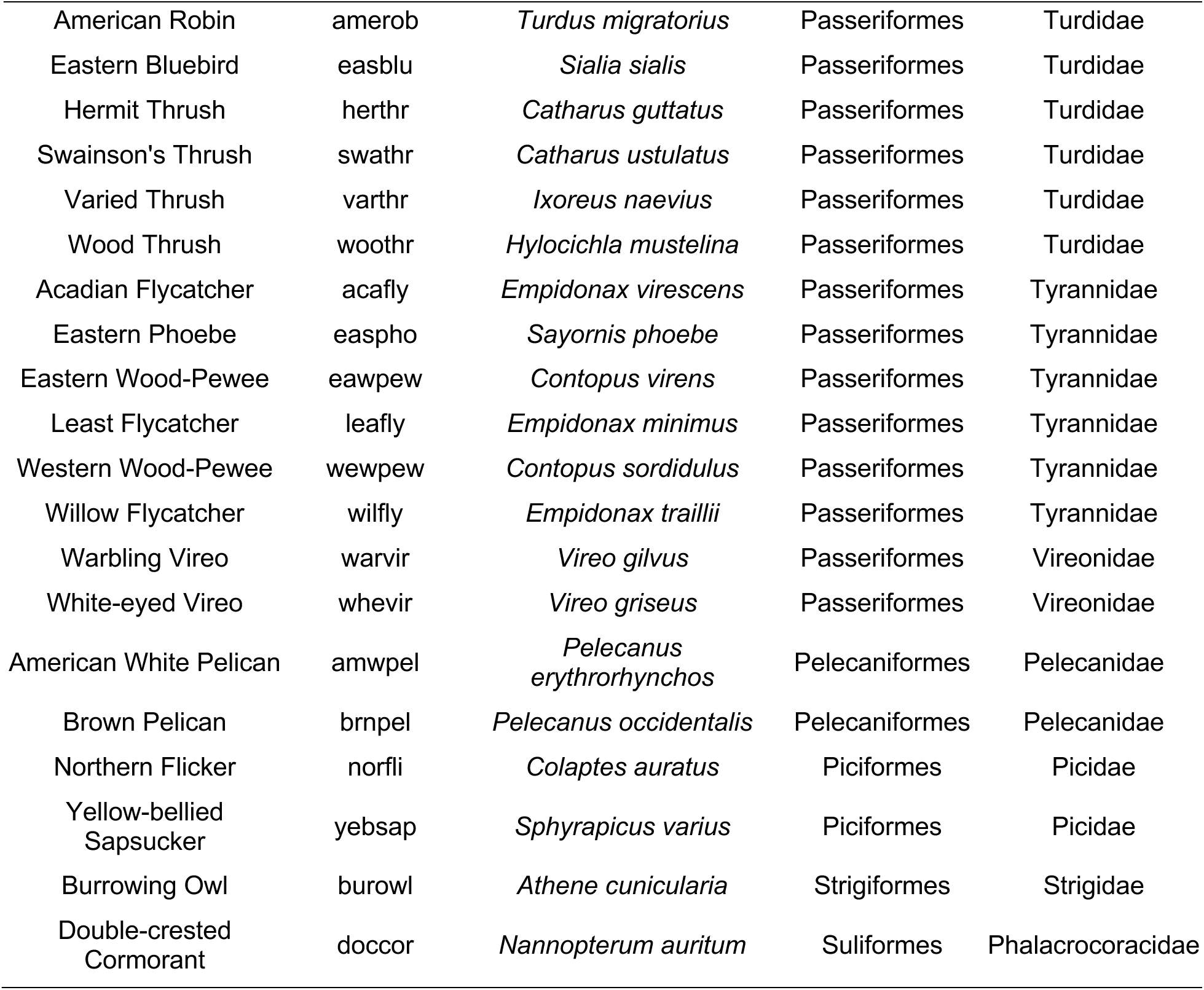
Species list for the 153 modeled species.

**Table S3.** Summarizing the biological plausibility of BirdFlow-simulated routes. A biologically plausible model should expect around 95% of the empirically observed biological metrics to occur within the 95% simulated intervals by BirdFlow models, and roughly 50% of the empirically observed biological metrics to occur within the 50% simulated intervals. This result indicates that BirdFlow-generated routes are biologically realistic and reflect population-level variation. More than 50% of observations occurred within the 50% simulated intervals for the number of stopovers, likely because the coarse weekly temporal resolution increases variability and reduces informativeness of stopover calculations. Stationary tracks were not included for straightness validation.

| Metric | Species | # GPS tracks for validation | % of empirical observations fall into the 95% simulated intervals | % of empirical observations fall into the 50% simulated intervals |
| --- | --- | --- | --- | --- |
| Straightness | American Woodcock | 452 | 95.58% | 61.73% |
|  | Blue-winged Teal | 63 | 63.49% | 26.98% |
|  | Broad-winged Hawk | 15 | 100.00% | 60.00% |
|  | Long-billed Curlew | 280 | 98.21% | 63.93% |
|  | Ovenbird | 9 | 88.89% | 11.11% |
|  | Turkey Vulture | 97 | 91.75% | 55.67% |
|  | Wood Thrush | 11 | 90.91% | 63.64% |
|  | **Averaged** |  | **89.83%** | **49.01%** |
| # stopovers | American Woodcock | 651 | 100.00% | 99.85% |
|  | Blue-winged Teal | 70 | 97.14% | 94.29% |
|  | Broad-winged Hawk | 19 | 100.00% | 100.00% |
|  | Long-billed Curlew | 296 | 100.00% | 100.00% |
|  | Ovenbird | 9 | 100.00% | 100.00% |
|  | Turkey Vulture | 118 | 100.00% | 94.92% |
|  | Wood Thrush | 11 | 100.00% | 100.00% |
|  | **Averaged** |  | **99.59%** | **98.44%** |
| Migration speed | American Woodcock | 651 | 94.93% | 56.22% |
|  | Blue-winged Teal | 70 | 74.29% | 31.43% |
|  | Broad-winged Hawk | 19 | 100.00% | 57.89% |
|  | Long-billed Curlew | 296 | 99.66% | 73.65% |
|  | Ovenbird | 9 | 55.56% | 22.22% |
|  | Turkey Vulture | 118 | 85.59% | 36.44% |
|  | Wood Thrush | 11 | 90.91% | 27.27% |
|  | **Averaged** |  | **85.85%** | **43.59%** |

## Supplementary methods

### Motus data preprocessing

Raw Motus data were summarized into “visits” for each time a bird visited a Motus station, with arrival and departure times for that event. These data were then validated using multiple criteria so that tracks generated from these summaries had reasonable estimated flight speeds, fell within expected species ranges, and there were at least 3 consecutive detections for any given visit. In addition, Motus stations which were deemed ‘too noisy’ were eliminated from the results to reduce the risk of introducing false positives.

### Transition sampling

We applied an iterative transition sampling procedure that considers the trade-off of computational cost and model performance improvement.

1. First, we iteratively sampled a single one-week transition from each of the tracks without replacement, until the maximum transition count was met or the target tracks run out of transitions to sample (or without any one-week transition to sample, as often seen in banding data), e.g., (𝑡_*_, 𝑥_*_) to (𝑡_G_, 𝑥_G_), and (𝑡_G_, 𝑥_G_) to (𝑡_H_, 𝑥_H_), where the 𝑡 is the weekly timestep and 𝑥 is the location corresponding to the timestep.
2. Second, if the size of the one-week transition set did not exceed the maximum transition count, we iteratively randomly sampled one 𝑛-timestep transition (𝑛 can be any positive integer from 2 to maximum week duration for the track) from each track without replacement, until the maximum transition count was met or the target tracks run out of transitions to sample, e.g., (𝑡_*_, 𝑥_*_) to (𝑡_I_, 𝑥_I_).

The rationale behind this sampling strategy is to prioritize the information-rich one-week transitions, while also integrating the information in longer transitions to tune and evaluate the prediction performance of BirdFlow models across seasons.

To constrain the transitions in a reasonable spatiotemporal range, only transitions that span less than 270 days with at least one end falling into migration seasons were sampled. We did not sample transitions with both ends falling into breeding or non-breeding seasons as we focused on the migratory periods, and since the species detectability can change vastly in stationary periods (especially during breeding and molting periods), which may cause spatiotemporal variation in S&T abundance surfaces that do not reflect movement (e.g., Socolar et al. 2025)

### Hyperparameter grid search settings

We reparametrized the hyperparameters for easier interpretation, defining the relative proportion of the observation term 𝜌 = 1/(1 + + 𝛽) and the relative proportion of the distance and entropy terms 𝜙 = 𝛽/𝛼, which allows us to rewrite Eq. 1 as

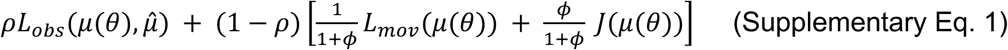

Since different combinations of parameters have different emphases on matching S&T abundance surfaces and regulating the route behavior patterns, this reparameterization allowed us to directly vary the overall emphasis of the observation weight with 𝜌 and vary the degree of stochasticity with 𝜙. In our grid search, we varied 𝜌 with values (0.95, 0.975, 0.99, 0.999, 0.9999) to emphasize the high importance of matching S&T abundance surface, 𝜙 across values (2, 4, 6, 8, 10), and 𝜖 with values (0.1, 0.2, 0.3, 0.4, 0.5, 0.6, 0.7, 0.8, 0.9). We chose these grid search values based on the empirical investigation in Fuentes et al. 2023 where none of the models with these parameter combinations perform the best with the extremal grid search values and the performance appears to vary smoothly as the hyperparameters change, which indicates a potentially optimal model enclosed in this hyperparameter surface. We configured parameters and passed them to the BirdFlowPy package (Fuentes 2022, Fuentes et al. 2023) for model fitting.

### Model decay analysis

The exponential regression can be formulated as:

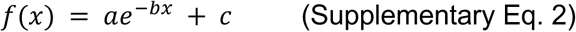

Where 𝑎, 𝑏, and 𝑐 are parameters and 𝑥 is the independent variable (time or distance horizon). The multi-species median regression line is calculated by taking the median value of each parameter across species, respectively. The maximum functional distance horizon is calculated as the value on the x-axis where the regression line intersects with a threshold, which was set to 0.05 for both the relative distance gain and the log-likelihood improvement (supplementary Fig. 4). For relative distance gain, a 0.05 value represents a horizon where the BirdFlow prediction is only 5% closer to the empirical resighting locations than the S&T abundance surfaces, and for log-likelihood improvement, 0.05 means the BirdFlow prediction is 1.05 times more likely to recover the empirical observation than a random sample from the S&T abundance surfaces. While these thresholds are arbitrary, we believe they generally reflect a functional horizon, beyond which the BirdFlow model is no longer meaningfully better than the original S&T prediction.

### Distributional summaries for each species

We calculated distributional summaries as explanatory variables for the best hyperparameters, including range size, spatial variation of abundance, the minimum, maximum, and centroid of longitude and latitude for different seasons. First, the eBird S&T abundance map was aggregated to 150km resolution and split into pre-breeding season, breeding season, post-breeding season, and non-breeding seasons. The season partitioning is queried from the eBird S&T dataset. Second, we averaged the abundance estimation across weeks for each spatial pixel by each season. Next, for each season, we calculated:

- Range size, as the maximum number of spatial pixels with abundance>0.
- Range abundance variation, as the standard deviation of the average-abundance variation across spatial pixels.
- Maximum and minimum range longitude and latitude: The maximum and minimum of the longitude and latitude across spatial pixels with abundance>0.
- Centroid of longitude and latitude: The abundance-weighted average longitude and latitude.

